# Evidence supporting an antimicrobial origin of targeting peptides to endosymbiotic organelles

**DOI:** 10.1101/2020.03.04.974964

**Authors:** Clotilde Garrido, Oliver D. Caspari, Yves Choquet, Francis-André Wollman, Ingrid Lafontaine

## Abstract

Mitochondria and chloroplasts emerged from primary endosymbiosis. Most proteins of the endosymbiont were subsequently expressed in the nucleo-cytosol of the host and organelle-targeted via the acquisition of N-terminal presequences, whose evolutionary origin remains enigmatic. Using a quantitative assessment of their physico-chemical properties, we show that organelle targeting peptides, which are distinct from signal peptides targeting other subcellular compartments, group with a subset of antimicrobial peptides. We demonstrate that extant antimicrobial peptides target a fluorescent reporter to either the mitochondria or the chloroplast in the green alga *Chlamydomonas reinhardtii* and, conversely, that extant targeting peptides still display antimicrobial activity. Thus, we provide strong computational and functional evidence for an evolutionary link between organelle-targeting and antimicrobial peptides. Our results support the view that resistance of bacterial progenitors of organelles to the attack of host antimicrobial peptides has been instrumental in eukaryogenesis and in emergence of photosynthetic eukaryotes.

## Introduction

Mitochondria and chloroplasts are eukaryotic organelles that evolved from bacterial ancestors through endosymbiosis (see [1] and [2] for recent reviews). These endosymbiotic events were accompanied by a massive transfer of genetic material from the bacterial ancestors to the host genome through what is known as Endosymbiotic Gene Transfer (EGT; [3]). Thus, to be successful, primary endosymbiosis required the establishment of efficient protein import machineries in the envelope membranes of the proto-organelle to re-import the products of the genes transferred to the nuclear genome. As a result, most mitochondrial and chloroplast genomes encode less than 100 proteins and the majority of proteins localised therein (ca. 1000 in mitochondria and >2000 in the chloroplast) are now translated in the cytosol and imported into the organelle [4,5]. Most nuclear-encoded proteins found in organelles harbour a targeting peptide (TP), an N-terminal presequence functioning as an address tag, *i.e.* determining the subcellular localisation of cargo proteins within endosymbiotic organelles [6]. TPs are recognised by the main mitochondrial and chloroplast translocation pathways [7,8] and destroyed upon import into organelles [4,5]. The emergence of TP-based import, despite being a key innovation enabling endosymbiosis and eukaryotism, remains poorly understood [8–10].

As described in a proposed scenario summarised in Supplementary Figure 1 for the emergence of the endosymbiotic protein import system [11], TPs may originate from antimicrobial peptides (AMPs). Archaea, bacteria and eukaryotes alike use antimicrobial peptides (AMPs) as part of their innate immune system to kill microbes, typically via membrane permeabilisation [12,13]. Numerous studies have established that AMPs consistently play a role in most symbiotic interactions [14,15], which argues for their involvement in the initial relationship between a host and a proto-endosymbiont. Extant heterotrophic protists employ AMPs to kill engulfed prey, which suggests that early eukaryotes likely used AMPs in a similar way against their cyanobacterial prey that ultimately became the chloroplast [16]. Similarly, the archaeal host will have delivered AMPs against the *α*-proteobacterial ancestor of mitochondria, whether it was a prey or an intracellular pathogen akin to Rickettsiales [17,18]. AMPs might also have been instrumental when considering mutualism at the origin of endosymbiosis, since present-day hosts use non-lethal concentrations of AMPs to control the growth of symbionts and to facilitate metabolic integration by enabling nutrient exchange [12,19].

Studies of the various extant strategies for microbial defence against AMPs have revealed instances where AMPs are imported into bacterial cells via dedicated transporters and then degraded by cytoplasmic peptidases, hereafter referred to as an “import-and-destroy” mechanism [20–26], which is strikingly reminiscent of TP-based import.

EGT thus would have started with incorporation of DNA fragments from lysed bacteria into the host genome, as observed in extant phagotrophic protists [27,28]. Endosymbiotic integration was then promoted through acquisition, by the proto-endosymbiotic bacteria, of an import-and-destroy mechanism to resist the AMP attack from the host. The serendipitous insertion of a bacterial gene downstream of an AMP coding sequence in the host genome, whether right upon EGT or after chromosomal rearrangements, then allowed the import of its gene product back into the protoorganelle via the very same inner membrane transporter that allowed the AMP-resistant endosymbiont to detoxify attacking peptides.

Indeed, extant TPs continue to show structural similarities to a type of AMP called Helical Amphiphilic Ribosomally-synthesised AMPs (HA-RAMPs), which are characterised by the presence of a cationic, amphiphilic α-helix [11,29,30]. Mitochondrial TPs (mTPs) contain a similar positively charged helix, the amphiphilic character of which is crucial for import [31–33]. The secondary structure of chloroplast TPs (cTPs) has been a matter of debate. They are longer than mTPs and contain parts that are not helical, such as an uncharged N-terminus thought to play a role in defining organelle specificity [6,34–36]. Most cTPs appear unstructured in aqueous solution [37], yet NMR studies using membrane-mimetic environments have demonstrated that cTPs also contain positively charged, amphiphilic α-helical stretches [38–40], suggesting cTPs fold upon contact with the chloroplast membrane [41].

Here, we tested the hypothesis that TPs have originated from host-delivered AMPs. If their common origin holds true, enduring similarities in their physico-chemical properties should have remain despite their large evolutionary distance. In addition, at least a subset of TPs and HA-RAMPs may still display some dual antimicrobial and organelle targeting activities. Here, we used the unicellular green alga *Chlamydomonas reinhardtii* to show that these two predictions are fulfilled, thus providing solid evidence that bacterial resistance to the host stands at the core of the emergence of eukaryotism.

We first assess in depth the physico-chemical properties of the various families of HA-RAMPs using a consistent set of descriptors and propose a more robust classification of these peptides compared to the current AMP families. We next provide computational evidence for the extensive overlap of the physico-chemical properties of TPs with those of a cluster of HA-RAMPs. Finally, we demonstrate experimentally that extant antimicrobial peptides are able to target a fluorescent reporter to either the mitochondria or the chloroplast of the *C. reinhardtii* and show that targeting peptides still display antimicrobial activity.

## Methods and Materials

### Sequence Data Set

The groups of peptides used in this study and the corresponding data sources are given in Table S1. Detailed information for each targeting, signaling, antimicrobial and random peptides are given in Tables S2 to S5. Antimicrobial peptides were extracted from the CAMP_*R*3_ database [42]. We selected the families of antimicrobial peptides based on the following criterion: i) activity experimentally validated, ii) documented amphiphilic a-helical structure, with at least one family member with a resolved 3D structure available in the Protein Data Bank [43], iii) documented activity of bacterial membrane destabilisation. For 3 families comprising peptides with very different structures, additional filtering was applied. We recovered only the bacteriocin of type IIa that are characterised by an amphiphilic helix [44] in BACTIBASE [45]. We selected only those defensins with a resolved 3D structure with at least 5 consecutive residues in a a-helix and only those cathelicidins with a resolved structure or defined as an amphiphilic helical peptide in [46] via UNIPROT. As a negative control, we retrieved the cyclotides of globular structure. TPs with experimentally-confirmed cleavage sites were recovered from proteomic studies (see Table S1). 200 eSPs were randomly extracted among 4707 confirmed eSPs from the Signal Peptide Website (http://www.signalpeptide.de). eSPs were selected so as to follow the length distributions of the peptides from HA-RAMP class I families. 200 random peptides were generated following the amino-acid frequencies observed in the Uniprot database and the length distribution of TPs. Data sets were curated to exclude sequences shorter than 12 amino acids or longer than 100 amino acids, following the length distributions of the peptides from HA-RAMP class I families.

### Peptide description and Auto-Cross Covariance (ACC) Terms

As amino acid descriptors, we used the Z-scales established by Hellberg and colleagues [47]. ACC terms between Z-scales were computed as described previously in [48]. ACC terms combine auto-covariance (same z-scale i = j) and cross-covariance (different z-scale i *≠* j) of neighboring amino acids over a window of 4 residues (lags (l) ranging from 1 to 4). There are thus nine nearest neighbor ACC terms for the 3 z-scales factors, yielding 36 ACC terms per peptide of length N. Each ACC term is defined for a given Z-scale couple (i,j) as follows:

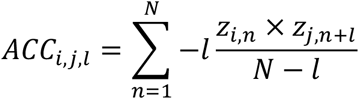

The Z-scales values were retrieved from the AAindex database (https://www.genome.jp/aaindex/) via the R package protr (v1.5-0). ACC terms were calculated with the acc function from the same R package and pre-processed by mean-centering and scaling to unit variance.

To assign a hydrophobicity value to a given peptide, we select the highest value obtained for a sliding window of 9 residues along the peptide. We used the hydrophobicity indices of amino acids estimated by octanol/water partitioning [49] to determine the mean hydrophobicity of the 9-residues window. The net charge of a peptide is the sum of the positively charged residues (arginine and lysine) and of the negatively charged residues (glutamate and aspartate) at pH 7.4.

The number of residues along the peptide that can theoretically adopt an amphiphilic helical structure is calculated as follows: The peptide is drawn along an a helical wheel and the longest region (of at least 9 residues) of the peptide that can adopt an amphiphilic helix is searched following the same criterion as in Heliquest [50]. The helix net charge corresponds to the net charge of these predicted amphiphilic helix.

### K-means clustering

Peptides were clustered based on the Euclidean distance defined above by k-means (scikit-learn Python package version 0.21.2). Centroid initialisation was performed with the ‘k-means++’ method with the best inertia among 100 runs for k ranging from 2 to 10. The selected k value (2) is that leading to the best average silhouettes coefficient -a measure of the clustering quality-[51] over all peptides.

### Distance trees

Euclidean distances between pairs of 36-dimensional vectors defining each peptide with their ACC terms were used to compute a distance tree between the studied HA-RAMPs, with the neighbor-joining implementation of the scikit-bio Python library, version 0.5.5. To evaluate the robustness of bipartitions on that NJ tree, we built 1000 trees from bootstrap ACC vectors and determined internode certainty (IC) and tree certainty (TC) measures [52] implemented in RaxML v8.2.12 [53]. Tree annotation and display was performed with iTOL v5.5 [54].

### Vizualisation of peptide properties

The two (or three) principal components of a Principal Component Analysis (PCA) of the peptides defined by their 36 ACC terms were used for visualisation of the peptide’s properties. The weights of each variable in the PCA are summarised by correlation circles in Supplementary Figure 5. Analyses were performed with the scikit-learn Python package version 0.21.2. Box plots were generated with the R ggplot2 package version 3.1.0.

### Detection of cTP motifs in HA-RAMPs

Scripts for finding Hsp70 binding sites and FGLK motifs were developed in R, exactly following the rules described by [55] and [56], respectively.

### Strains and culture conditions

*C. reinhardtii* cells derived from wild-type strain T222+ *(nit1, nit2, mt+)* were grown in mixotrophic conditions in Tris-acetate-phosphate (TAP) medium [57] under ~30 μmol photons m^-2^ s^-1^ at 25°C, either in 200 μL in 96-well plates for 3-4 days or in agitated Erlenmeyer flasks of 200 mL. Constructs were transformed into strain T222+ by electroporation and transformants, selected for paromomycin resistance, were screened for high Venus expression in a fluorescence plate reader (CLARIOstar, BMG labtech) as described in [58].

### Generation of constructs

Constructs were made by inserting sequences coding for candidate peptides directly upstream of the Venus start codon in plasmid pMO611, kindly provided by the Pringle lab. pMO611 is a derivative of the published bicistronic expression plasmid pMO449 [58] in which translation of the eight first RBCS codons ahead of the Venus coding sequence was prevented by mutating the start codon to CTG. To increase expression of the AMP constructs, we modified plasmid pMO611 further by inserting RBCS2 intron 2 within Venus, 73 bases downstream of the initiation codon. Introns do not influence targeting. Native Chlamydomonas TP sequences were amplified from strain T222+ genomic DNA, while codon optimised AMP gene sequences were synthesised by Eurofins Genomics. Sequences used are detailed in Table S6. Peptide constructs were assembled and integrated upstream of Venus using the NEBuilder HiFi Assembly kit (New England Biolabs). Correct assembly was verified by sequencing of inserts and flanking regions. Linear transformation cassettes were excised from plasmids with *EcoRV* (New England Biolabs) prior to transformation.

### Microscopy

For confocal imaging (Figure 5 and Supplementary Figures 9), cells were subjected to 0.1 *μM* MitoTracker Red CMXRos (ThermoFisher) for 30 minutes in the dark and washed with TAP prior to imaging on an upright SP5 confocal microscope (Leica). Venus (excitation 514 nm/532-555 nm emission) and MitoTracker (561 nm/573-637 nm) were imaged sequentially to avoid crosstalk, each alongside chlorophyll autofluorescence (670-750nm emission) to eliminate cells that had moved between images (chlorophyll data from 514 nm excitation shown in figures). Epifluorescence images (Supplementary Figures 7 and 10) were taken on an Axio Observer.Z1 inverted microscope (Zeiss) equipped with an ORCA-flash4.0 digital camera (Hamamatsu) and a Colibri.2 LED system (Zeiss) for excitation at 505 nm for Venus (filter 46HE YFP shift free, 520-550nm emission) and 470 nm for chlorophyll autofluorescence (filter set 50, 665-715nm emission) with cells in poly-L-lysine (Sigma Aldrich) coated 8-well μ-slides (Ibidi). Image brightness was adjusted for presentation in figures and cyan, yellow and magenta linear lookup tables, assigned to MitoTracker, Venus and chlorophyll channels respectively, in Fiji (http://fiji.sc/Fiji, version 2.0.0-rc-69/1.52p). To quantify co-localisation (Supplementary Figure 9), cells were cropped out of larger fields-of-view using a standard region-of-interest quadratic box with 13 *μm* side length in Fiji. Fiji was also used to measure image background intensities from outside the cell for the Venus channel, or from within the cell but outside the organelle in the case of the MitoTracker and chlorophyll channels. Backgrounds were then substracted and Pearson correlation coefficients were calculated as a measure of co-localisation [59].

### Biochemistry

Chloroplasts and mitochondria were isolated essentially as described in [60], with the following modifications. Protease assay data (Figure 6C,D) and purity data (Figure 6A,B) came from two different extractions (a) and (b) respectively, which were done as follows. 2 L cultures inoculated either into TAP 48h before the experiment and grown at 50 μE m^-2^ s^-1^ (a), or in Minimal Medium 72h before the experiment and grown aerated under at 200 μE m^-2^ s^-1^ (b), were kept in darkness the night preceding the experiment. Cells were washed twice in 20 mM HEPES-KOH pH 7.2 (a), or washed once in 10 mM HEPES-KOH pH 7.2 and then subjected to an acid shock to remove flagella as described in Craige et al. (2013) (1min at pH 4.5 by adding 1M acetic acid, then neutralising back to pH 7.2 with 1M KOH) (b). Working in a cold room, cells were then resuspended in 50 mL ice cold breaking buffer (300 mM Sorbitol, 3 mM MgCl2, 5 mM EDTA, 0.5% Polyvinylpyrrolidone PVP40, in 67 mM HEPES-KOH pH 7.2) and disrupted by nebulisation with Argon using either one pass at 40 psi (a), or four passes at 80 psi (b), after which an aliquot was set aside. A brief centrifugation up to 4000 g (8 °C, manual stop when reaching 6000 RPM, Beckman JA-12) was used to separate chloroplast and mitochondrial fractions in pellet and supernatant respectively. Chloroplasts were recovered at either the 45/70% (a) or the 20/40% (b) interphase of a 12 ml discontinuous Percoll gradient (in chloroplast wash buffer: 300 mM Sorbitol, 3 mM MgCl2, in 67 mM HEPES-KOH pH 7.2) after 2 h of ultracentrifugation (4 °C, 5000 RPM, Beckman SW41, slow acceleration and deceleration) and washed in 5 volumes of chloroplast wash buffer (8 °C, manual stop when reaching 3000 RPM, Beckman JA-20, acceleration: 5, deceleration: 0). Mitochondria were purified using a series of differential centrifugations: 20 min at 1500 g to pellet residual chloroplasts (8 °C, 3500 RPM, Beckman JA-12), 20 min at 20 000 g to pellet mitochondria (8 °C, 11 300 RPM, Beckman JA-13.1), 40 min at 20 000 g (8 °C, 11 300 RPM, Beckman JA-13.1) to run a 30 ml 20% Percoll purification (in 250 mM Sorbitol, 1 mM EDTA, 0.5% PVP40, in 10 mM MOPS-KOH pH 7.2) of which the bottom ~3 mL were kept, and further centrifuged 20 min at 20 000 g (8 °C, 11 300 RPM, Beckman JA-13.1) to pellet mitochondria after Percoll removal by dilution with 40 mL mitochondrial wash buffer (250 mM Sorbitol, 1mM EDTA, in 10 mM phosphate buffer pH 7.2). Isolated organelle pellets were resuspended in 100 μL SEM (250 mM Sucrose, 1 mM EDTA, in 10 mM MOPS-KOH pH 7.2). For the proteinase assay [60], a 22 μL aliquot of isolated organelles was treated with 1.1 μL 25 % Triton X-100 for 5 min on ice, another with an equal volume of water. Each sample was split into two aliquots of 10.5 μL, one of which was treated with 0.5 μL proteinase K 20x stock in SEM (final concentration: 150 *μ*g/mL) for 15 min on ice, and the other with an equal volume of SEM. All aliquots were then treated with 0.25 μL 100 mM PMSF and 2.8 μL 5x storage buffer (5x Roche cOmplete™ Mini proteinase inhibitor cocktail, 50 mM NaF, 1 M DTT, 1 M Na2CO3) and stored at −20°C. Of each isolated organelle and whole cell sample, an aliquot was precipitated overnight at 4°C by adding 900 μL 80% Acetone, pelleted for 10min (4°C, max speed, Eppendorf 5415 D benchtop centrifuge), dried for ~10min under vacuum, resuspended in protein resuspension buffer (2% SDS, 10 mM NaF, 1x Roche cOmplete Mini proteinase inhibitor cocktail, in 100 mM Na2CO3) and used for protein quantification by BCA assay (ThermoFisher Scientific). Samples were then run on either 4-15% (a) or 8-16% (b) precast gels (Biorad), transferred onto 0.1μm nitrocellulose membranes and used for immunoblot detection. To be able to probe a very limited amount of sample with multiple antibodies, membranes were cut horizontally into separately treated strips. Primary antibodies raised against the following proteins were used in block buffer (5% BSA, 0.1% Tween-20, in PBS) at the indicated dilutions: α-Tubulin (Sigma Aldrich T5168, 1:50,000), FLAG (Agrisera AS15 2871, 1:10,000), COXIIb (Agrisera AS06 151, 1:10,000), BIP (Agrisera AS09 481, 1:2000), CF1β (homemade, rabbit, 1:50,000) [62], OEE2 (homemade, rabbit, 1:2000) [63], RBCS (kindly supplied by Spencer Whitney, rabbit, 1:20,000) [64], NAB1 (Agrisera AS08 333, 1:10,000). ECL signals were recorded on a ChemiDoc Touch (Biorad). Blots were cropped and final figures assembled in Powerpoint.

### Antimicrobial activity assays

Standard minimum inhibitory concentration broth microdilution assays in the presence of BSA/acetic acid were performed in triplicate as described in [65] using peptides chemically synthesised to ≥95% purity (Proteogenix). Dilution series are based on net peptide content, calculated by multiplying the dry weight by %N and by purity. %N is a measure of the peptide (rather than salt) fraction in the lyophilised product, while purity, provided by the manufacturer, is the fraction of the peptide with the desired sequence among all supplied peptides. Peptide sequences are listed in Table S7.

### Statistical analysis

Chi^2^ tests were used to analyse the distribution of peptides according to their functional group among the different k-means clusters (Figure 1). Wilcoxon test for all paired comparaison with a Holm correction were used to analyse the distributions of peptide features (Figure 2 and Figure 3). One-way analysis of variance (ANOVA) and Tukey post-hoc were used to compare the Pearson correlation coefficients to analyse fluorescence intensities (Supplementary Figure 9). A p-value threshold of 0.05 was used for all tests. All statistical calculations were performed with the stats package (v3.6.2) of the R version 3.6.1 and with functions from the Python scipy module (v1.2.3).

**Figure 1.**
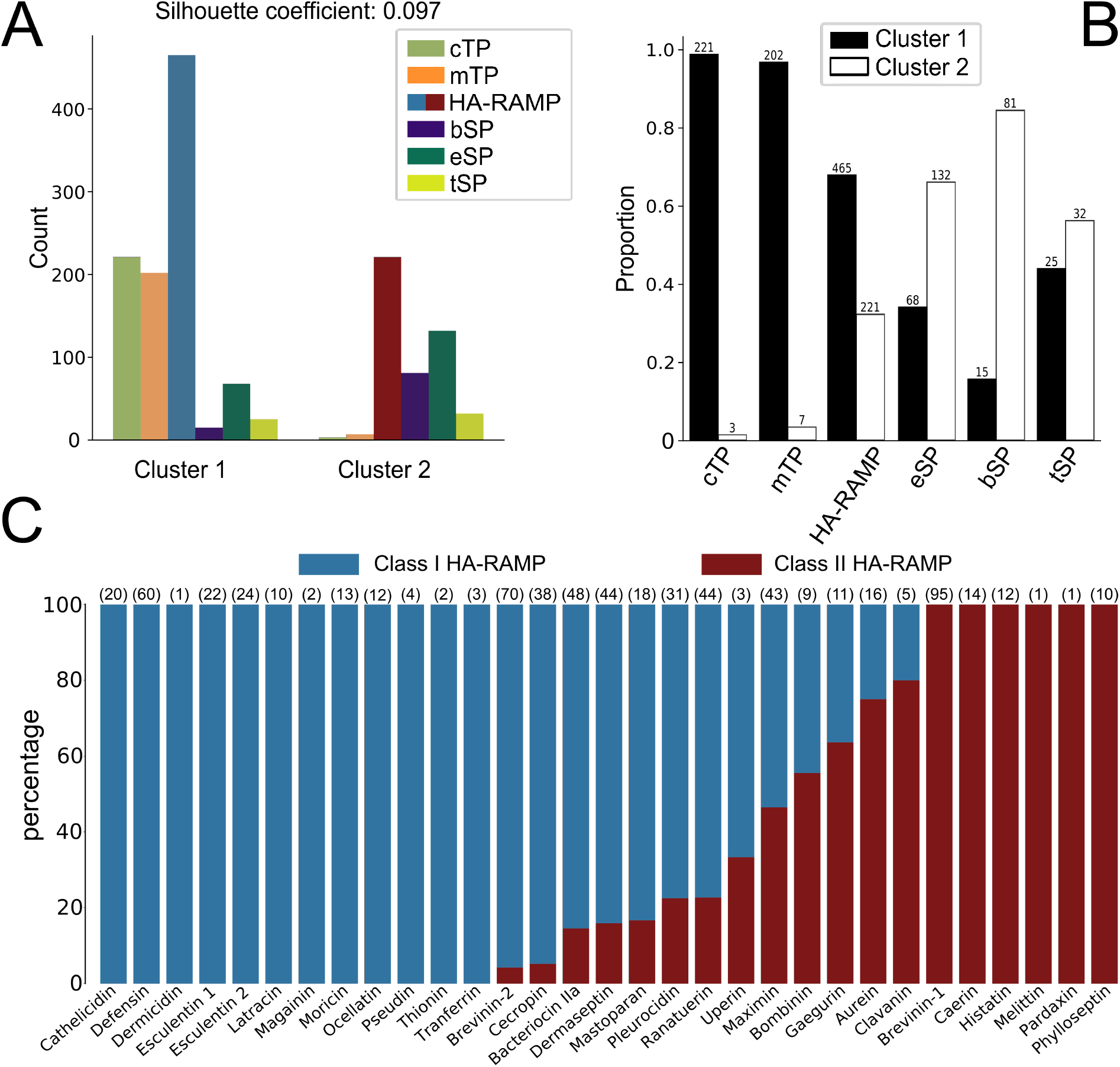
TPs cluster with certain types of HA-RAMPs. (A) Contribution of the different groups of peptides, as described by 36 ACC terms, to the 2 clusters obtained by k-means (Chi^2^ Pearson test, *p* < 4.94 · 10^-324^). The average silhouette coefficient is indicated above the graph. HA-RAMPs are depicted in blue in cluster 1 and in red in cluster 2. (B) Proportion of peptides among the 2 clusters for each group. Total number of peptides is indicated above bars. See figure panel for colour code. (C) Percentage of peptides from the HA-RAMP families described in the literature among classes I and II.

**Figure 2.**
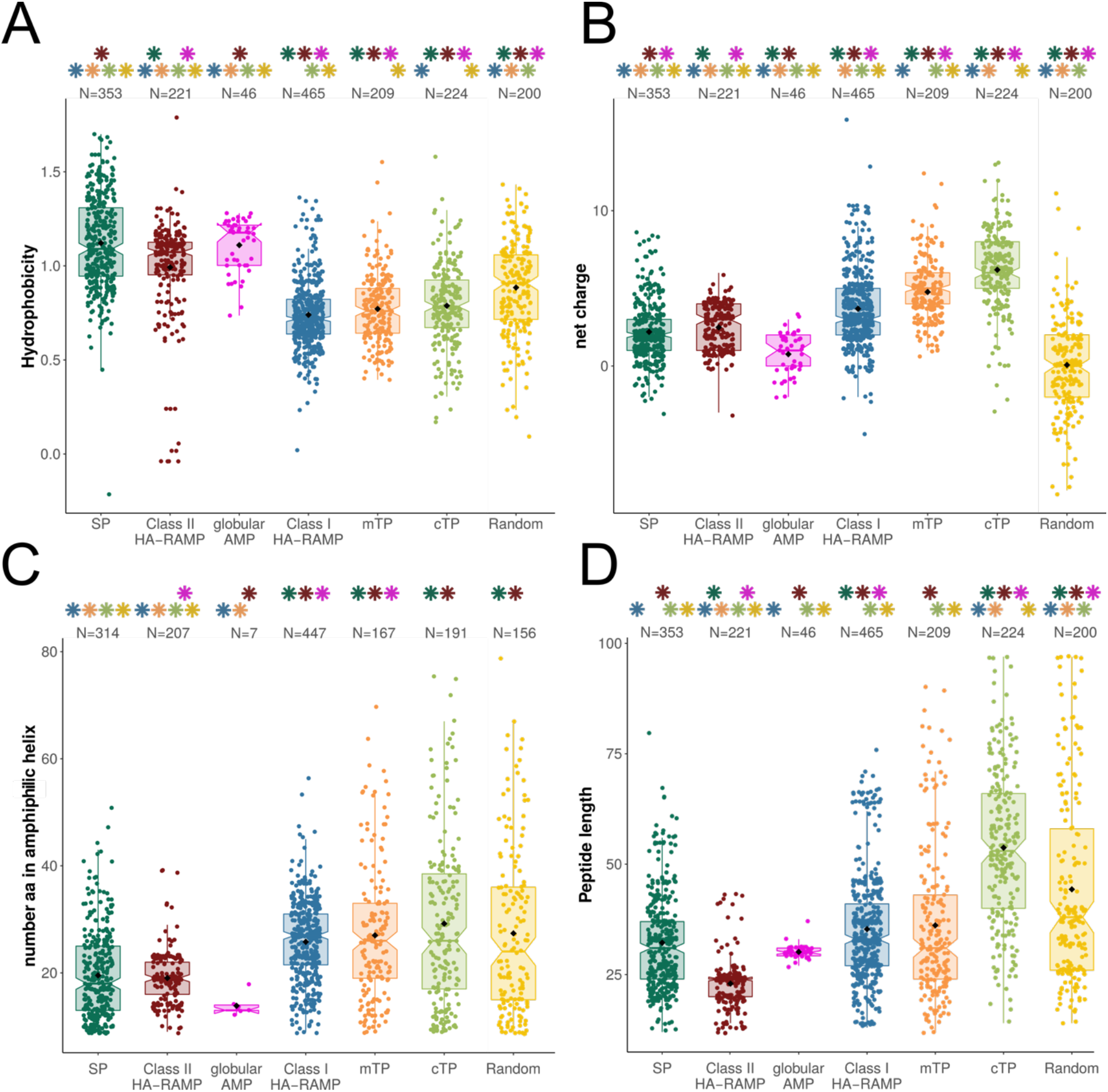
TPs and Class I HA-RAMPs display similar general features. Distributions are represented by coloured box plots and individual values by coloured points; black diamond indicate mean values (A) Maximum hydrophobicity for a 9-residues window along the peptide. (B) Net charge of the peptide. (C) Number of amino acids that can formally adopt an amphiphilic α helical structure within the peptide (minimum of 9 residues). (D) Peptide length in amino acids. Numbers above distributions indicate the number of peptides represented (note that some peptides have no predicted amphiphilic helix). Stars indicate significant differences (Wilcoxon tests, p-value < 0.05) between that distribution and the distribution with the same colour as the star.

**Figure 3.**
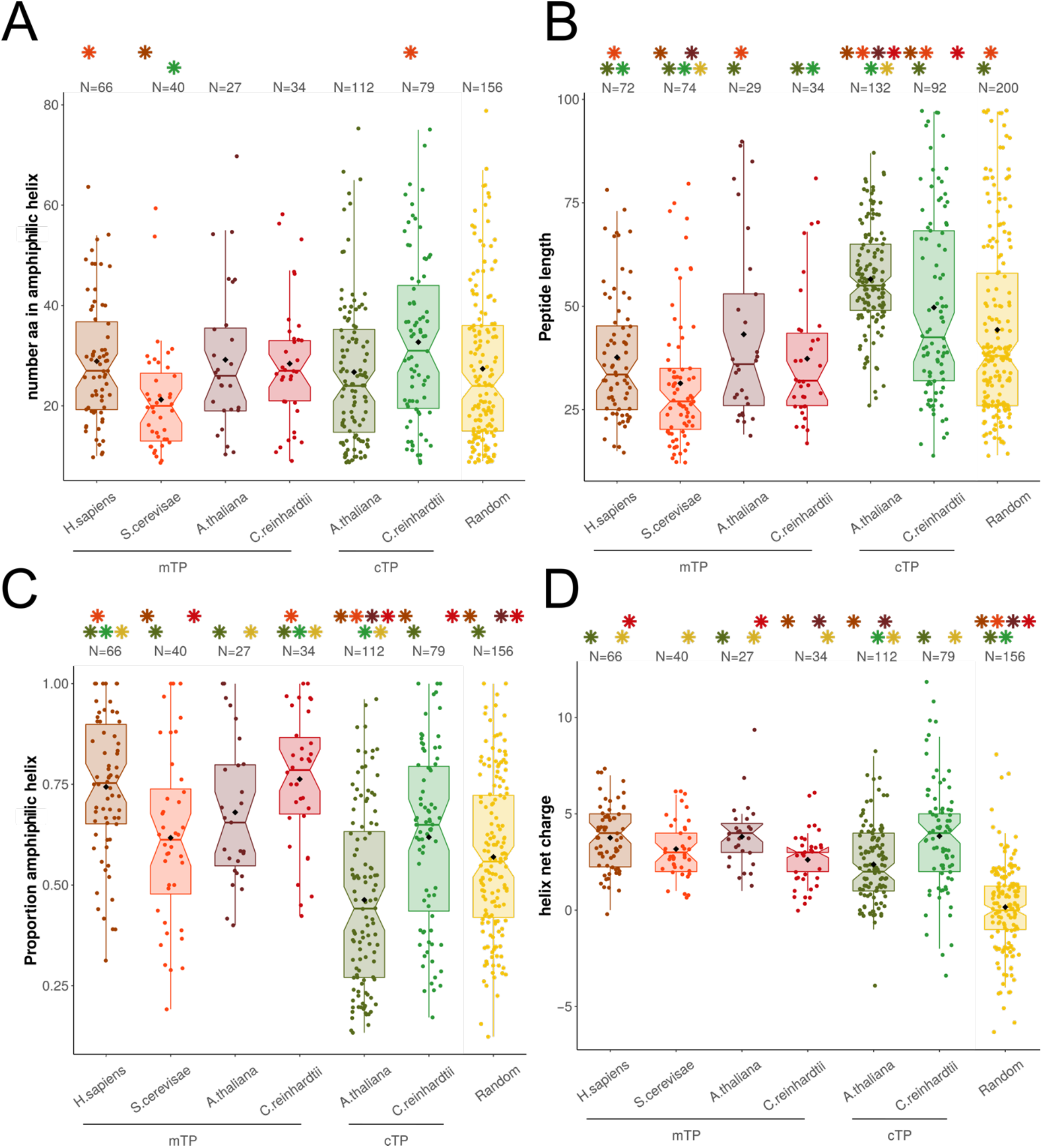
TPs display amphiphilic properties. Distributions are represented by coloured box plots as in Figure 2. (A) Number of amino acids that can formally adopt an amphiphilic α helical structure within the peptide. (B) Peptide length in amino acids. (C) Proportion of peptide predicted as amphiphilic (D) Net charge of the predicted helix. Numbers above distributions indicate the number of peptides represented (note that some peptides have no predicted amphipatic helix). Stars indicate significant differences (Wilcoxon tests, p-value < 0.05) between that distribution and the distribution with the same colour as the star.

### Code availability

All Python and R in-house scripts are available at https://github.com/Clotildegarrido/Peptides_Analysis

## Results

### TPs and HA-RAMPs share a common set of physico-chemical properties

#### Peptide families and their descriptors

We performed a comparative analysis of different functional groups of peptides (Supplementary Table S1). We selected TPs from *Chlamydomonas reinhardtii, Arabidopsis thaliana, Saccharomyces cerevisiae* and *Homo sapiens* for which both the subcellular location of the cargo protein and the cleavage site have been experimentally determined (Supplementary Table S2). We retrieved from the CAMP_*R*3_ database [42] 31 HA-RAMP families with a documented amphiphilic domain (see Material and Methods) and, as negative control, the cyclotide family of globular AMPs (Supplementary Table S4). We also considered a set of peptides, hereafter referred to as secretory signal peptides (SPs), which function as address tags, just as TPs do, but target a different subcellular compartment. SPs have a well-established evolutionary link and all use Sec-type translocation systems [66–68]. We retrieved as SPs the bacterial SPs (bSPs) that target proteins for periplasmic secretion, their eukaryotic relatives (eSPs) targeting proteins to the endoplasmic reticulum, and thylakoid SPs (tSPs), more commonly referred to as thylakoid transit peptides, that target proteins to the thylakoids [6,68] (Supplementary Table S3). Lastly, we generated a set of random peptides (Supplementary Table S5).

All peptides were less than 100 amino acids long. On average they are comprised of 45 residues for TPs, 32 residues for HA-RAMPs and 31 residues for SPs. Because TPs and HA-RAMPs are short peptides with very limited sequence similarity, classical phylogenetic inferences were not applicable [6]. Thus, to evaluate the likelihood of an evolutionary relationship between TPs and HA-RAMPs, we resorted to their physico-chemical properties rather than to their primary sequences and used the amino-acid descriptors ‘Z-scales’ defined by Hellberg [47]. Each of the 20 amino acids is therein described as a set of three values, which correspond to the first three linear combinations (principal components) of 29 physico-chemical properties measured experimentally. These three Z-scales reflect mostly hydrophobicity (z1), bulkiness of the side chain (z2) and electronic properties (z3). A comparative study of 13 types of amino acid descriptors showed that these three Z-scales are sufficient to explain the structure-activity variability of peptides [69,70]. To account for interdependencies between residues – i.e. the properties of the whole peptide – each peptide was defined by 36 terms corresponding to auto-cross covariances (ACC) between Z-scale values [48] within a 4-neighbor window, which mimics a single α-helix turn of 3.6 residues (see “Peptide description” in the Materials and Methods section).

#### HA-RAMPs can be divided into two distinct classes

Many HA-RAMPs families have been defined on a rather descriptive basis in the literature and the criteria used to group peptides differ from one family to another. To draw a more consistent picture of the diversity of HA-RAMPs, we performed a k-means clustering of our 686 selected HA-RAMPs, together with the 353 SPs and 433 TPs, based on the Euclidean distances between their 36 ACC vectors (Figure 1A). Among clustering with k varying from 2 to 10, clustering with k=2 gave the highest average of silhouette coefficients [51] for all peptides (reflecting the consistency of the clustering). HA-RAMPs distributed between the two clusters in a 2/3 vs. 1/3 proportion (Figure 1B). The 68% of HA-RAMPs that grouped in cluster 1 will be hereafter referred to as class I HA-RAMPs, whereas those in cluster 2 will be referred to as class II HA-RAMPs. 60% of the families described in the literature fitted well either into cluster 1 or cluster 2, which supports their classification as families of distinct physico-chemical properties (Figure 1C). However, 13 families contained peptides distributed in both clusters, calling for further investigation of their identification as members of a same family.

To get a more detailed picture of their similarity relationships, we performed a neighbour-joining (NJ) clustering of all HA-RAMPs based on the Euclidean distances between their 36 ACC vectors (Supplementary Figure 2). The most external bipartitions of the clustering tree are highly supported, while internal ones are less supported. Class I and class II HA-RAMPs are not intermingled on that tree and tend to form robust homogeneous sub-clusters. When considering peptides according to their antimicrobial families described in the literature, their distribution along the tree was patchy (outer colour circle on Supplementary Figure 2).

To better handle the differences between class I and class II HA-RAMPs, we compared their features in terms of length, hydrophobicity, net charge and number of residues that can theoretically adopt an amphiphilic helical structure (Figure 2). The two classes indeed had rather distinctive traits: compared to class II HA-RAMPs, class I HA-RAMPs have significantly lower hydrophobicity (Figure 2A), higher net charge (Figure 2B), longer amphiphilic helices (Figure 2C) and are overall longer peptides (Figure 2D).

#### TPs and class I HA-RAMPs share a set of physico-chemical properties

Based on the k-means classification of HA-RAMPs in two classes with distinct traits, we further investigated the properties of TPs and SPs relative to those of HA-RAMPs. Figure 1 shows that all TPs, but a few isolated ones, clustered with class I HA-RAMPs. The majority of SPs grouped together in the other cluster. The grouping of most TPs with a large subset of class IHA-RAMPs proved very robust, being maintained for k values increasing up to 10 (Supplementary Figure 3). Moreover, when grouped together, class I HA-RAMPs and TPs are always the most abundant peptides in the cluster (the left one in Supplementary Figure 3). In contrast, the grouping of class II HA-RAMPs and SPs vanishes with increasing k values, being lost for k values greater than 6 (Supplementary Figure 3). These observations reveal strong similarities among a large subset of class I HA-RAMPs and TPs, but not between SPs and class II HA-RAMPs (see below).

The basis for this distinct clustering is documented in Figure 2: TPs and class I HA-RAMPs follow the same trends, away from the more hydrophobic class II HA-RAMPs and from SPs that bear a well-documented hydrophobic stretch (Figure 2A). Furthermore, TPs and class IHA-RAMPs all form amphiphilic helices of similar length (Figure 2C). Interestingly, randomly generated peptides contain amphiphilic stretches of similar length, albeit without the characteristic cationic character of TPs and HA-RAMPs. By contrast, SPs and globular cyclotides contain significantly shorter amphiphilic stretches, suggesting amphiphilicity may be actively selected against. Shorter amphiphilic helices in class II HA-RAMPs are due to the shorter overall length of these peptides (Figure 2D). The fact that cTPs are significantly longer than class I HA-RAMPs and mTPs while containing amphiphilic stretches of similar length is in line with the idea that cTPs contain additional sequence elements [5,36]. The control group of globular cyclotides display the highest hydrophobicity (Figure 2A) and the shortest amphiphilic helices (Figure 2C), as expected from their globular nature.

Because mTPs are well recognized as being of amphiphilic nature whereas cTPs are often referred to as unstructured peptides, we carefully reassessed their amphiphilic properties in the various organisms that we used in the present study (Figure 3). On average, cTPs and mTPs form amphiphilic helices of similar length, but for *S. cerevisiae* mTPs which are much shorter (Figure 3A). Both mTPs and cTPs are longer in *A. thaliana* than in *C. reinhardtii*, which results in a higher proportion of amphiphilic sequence in mTPs and cTPs from the latter (Figure 3C), in line with previous reports that algal cTPs resemble plant mTPs [71]. However, irrespective of species, cTPs are longer than mTPs, thus displaying smaller proportion of amphiphilic sequence. Taken together, these characteristics explain why cTPs have been reported as less amphiphilic than mTPs, despite the presence of a *bona fide* amphiphilic helix. It is of note that the amphiphilic helices detected in random peptides have widely different characteristics since they also involve negatively charged residues which are largely excluded from those detected in TPs and HA-RAMPs (Figure 1C and 3D): the majority of the amphipathic helices are positively charged in TPs (92%), when they are only 47% of those identified among random peptides.

To better characterise the physico-chemical properties that are most discriminatory between SPs, HA-RAMPs and TPs, we performed a Principal Component Analysis (PCA) of these peptides described by their ACC vectors. Figure 4 presents a PCA without class II HA-RAMPs (see Supplementary Figure 4 for a PCA with class II HA-RAMPs). The separation between all peptides is provided by a combination of the contributions of various ACC terms to the two principal components (PC1 and PC2). As shown by the contributions of the various ACC terms (Supplementary Figure 5A), the terms reflecting the coupling between electronic and steric properties of the residues from the opposite faces of the amphiphilic helix are the main contributors to PC1 whereas the terms reflecting the hydrophobic and steric properties of the residues along the same face of the helix mostly contribute to PC2. When considering the amphiphilic helical domain of a peptide, these terms respectively reflect the electronic constraints that residues have to match on the same face of the a-helix and the amphiphilic constraints between residues from opposite faces.

**Figure 4.**
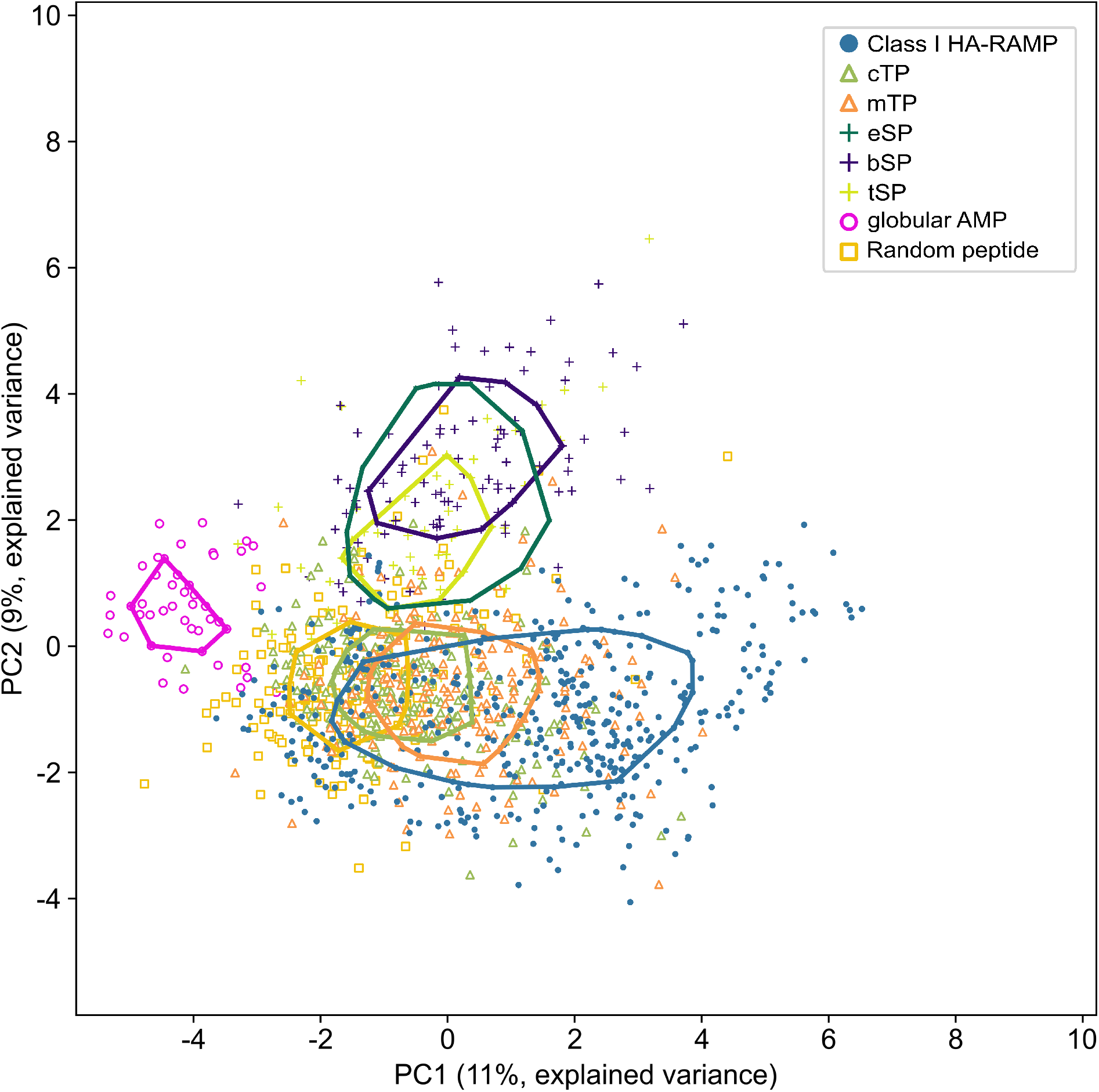
TPs and class I HA-RAMPs share similar physico-chemical properties. PCA is performed on peptides described by 36 ACC terms. Peptide positions are plotted along the first (PC1) and second (PC2) principal component on the x and y axes respectively, with explained variance in parenthesis. Solid lines represent the convex areas containing the 50% most central peptides of each group. Groups are class I HA-RAMPs (blue) as defined in Figure 2, TPs (mTPs, orange triangle; cTPs, green triangle), SPs (eSPs, dark green cross; bSPs, indigo cross; tSPs, light green cross) and control peptides (globular AMP pink circle; random peptide yellow square). See Supplementary Figure 5A for the contribution of ACC terms PC1 and PC2 and Supplementary Figure 8 for PCA with class II HA-RAMPs.

The evolutionarily-linked and hydrophobic tSPs, bSPs and eSPs co-localise on the top of the graph, away from TPs and class I HA-RAMPs (Figure 4). Class I HA-RAMPs occupy the bottom of the graph with an amphiphilic gradient from left to right, overlapping with TPs on the left side. TPs form a single overlapping spread, but mTPs show a tendency for higher values along PC1 than cTPs, in agreement with amphiphilic helices taking up a higher proportion of each peptide in mTPs. These observations are in line with k-means clustering (Figure 2, Supplementary Figure 3) where TPs group with class I HA-RAMPs, apart from SPs. As control groups, we display on Figure 4 the distribution of globular AMPs that occupy a separate part of the physico-chemical space to the left of the graph, reflecting their widely different structure, despite a shared antimicrobial function. The overlap of random peptides is higher with HA-RAMPs and TPs than with SPs. Note that part of this overlap originates from amphiphilic features of random peptides which are born by negatively charged residues, at variance with the positively charged amphiphilic helices present in TPs and Class I HA-RAMP (Figure 3D).

When considering all HA-RAMPs together with SPs and TPs in the plane defined by PC1 and PC2 (Supplementary Figure 4A), TPs are almost completely enclosed within the convex area of class I HA-RAMPs. The partial overlap of SPs with class II HA-RAMPs stems from their more hydrophobic character, compared to class I HA-RAMPs. By contrast, in the plane defined by PC1 and PC3 (reflecting the hydrophobic properties of the residues along the same side of the helix), class II HA-RAMPs group closer to class I HA-RAMPs and away from SPs (Supplementary Figure 4B) reflecting their different amphiphilic character (Supplementary Figure 4D). However, TPs still overlap with class I HA-RAMPs in the PC1/PC3 plane, reflecting a much tighter physico-chemical relatedness, as already observed when comparing the general features of the peptides (Figure 2) and within k-means clustering (Supplementary Figure 3).

### HA-RAMPs and TPs show dual targeting and antimicrobial activities

#### A TP cleavage-site fragment is required for import of the Venus reporter

To assess the targeting activity of AMPs we used a bicistronic expression system based on ribosome re-initiation as described by Onishi and Pringle [58] with coding sequences for candidate peptides inserted upstream of a Venus fluorescent reporter [72]. In this bicistronic system, the stop codon of the fluorescent reporter and the initiation codon of the selectable marker are separated by only six nucleotides (TAGCAT), which is sufficient to ensure robust expression of both genes in *C. reinhardtii*. Compared to classical expression systems where the selectable marker is driven by a separate promoter, bicistronic expression results in a much higher fraction of recovered transformants showing expression of the gene of interest [58].

Mitochondria and chloroplasts were imaged respectively using a MitoTracker dye and chlorophyll autofluorescence. In the absence of Venus (expression of the selectable marker only), some crosstalk is visible in the Venus channel (Supplementary Figure 6A), which appears to originate from thylakoid localised pigments and in particular from the eyespot. In the absence of a presequence (Supplementary Figure 6B) Venus remains cytosolic.

Surprisingly, the fluorescent reporter was equally cytosolic when the Rubisco activase cTP (RBCA-cTP) up to the cleavage site was included upstream of Venus (Supplementary Figure 6C). For import into the chloroplast, a stretch of 23 downstream residues was required (RBCA-cTP+), to reconstitute a native cleavage site (Supplementary Figure 6D). This finding is in line with previous efforts to target reporters to the chloroplast [73,74]) and led us to include residues −10 to +23 with respect to the cleavage site in subsequent constructs. This cleavage site fragment (RBCA-cs) by itself displayed no capacity for directing the Venus reporter into either organelle (Supplementary Figure 6E). We note that in addition to colocalising with chlorophyll, the Venus reporter driven by RBCA-cTP+ was abundant around or within the pyrenoid, the native location of RBCA (Supplementary Figure 6D). The pyrenoid is a proteinaceous structure of importance to the algal carbon-concentrating mechanism that contains a lower density of thylakoid membranes than the rest of the chloroplast. As a result, it is visible as a characteristic dark zone in chlorophyll auto-fluorescence at the apex of the chloroplast [75], making Venus accumulating at this site easy to spot. Mitochondrial localisation of Venus driven by a native *C. reinhardtii* mTP (CAG2-mTP+), including post-cleavage site residues, is characterised by a tell-tale pattern [75] and co-localisation with the MitoTracker signal (Supplementary Figure 6F). When the residues of this same mTP were rearranged so as to impede the formation of an amphiphilic helix, the resulting peptide was no longer able to target the reporter (Supplementary Figure 7A).

#### HA-RAMPs target Venus to endosymbiotic organelles

To assess the organelle targeting ability of HA-RAMPs, we selected five class I peptide candidates that clustered with TPs, by k-means clustering based on their Euclidean distances: bacillocin 1580 and enterocin HF from the bacteriocin IIA family, the cecropin sarcotoxin-1D, brevinin-2ISb from the brevinin-2 family, and the well-studied magainin II. These candidates localise next to TPs in our PCA analysis (Supplementary Figure 8A). When fused alongside RBCA-cs upstream of Venus and expressed in *C. reinhardtii* (Figure 5), both bacillocin 1580 (Figure 5A) and enterocin HF (Figure 5B) give rise to a fluorescence signal that is co-localised with chlorophyll auto-fluorescence, in line with their proximity to cTPs in the PCA (Supplementary Figure 8A). There is also a marked accumulation around the pyrenoid, particularly for bacillocin 1580. Although closer to mTPs in our PCA analysis (Supplementary Figure 8A), sarcotoxin-1D also targeted Venus to the chloroplast (Figure 5C), in line with the fact that some cTPs are found in the vicinity of our sarcotoxin-1D construct (Supplementary Figure 8A). Brevinin-2Isb on the other hand, proximal both to mTPs and cTPs in PCA, resulted in Venus fluorescence showing the typical pattern of mitochondrial localisation, co-localising with the MitoTracker dye (Figure 5D). Magainin II also targeted Venus to the mitochondria (Figure 5E), as might be expected from the construct most distal to cTPs in our PCA (Supplementary Figure 8A). Class I HA-RAMPs are thus capable of targeting a cargo protein to either type of endosymbiotic organelles.

**Figure 5.**
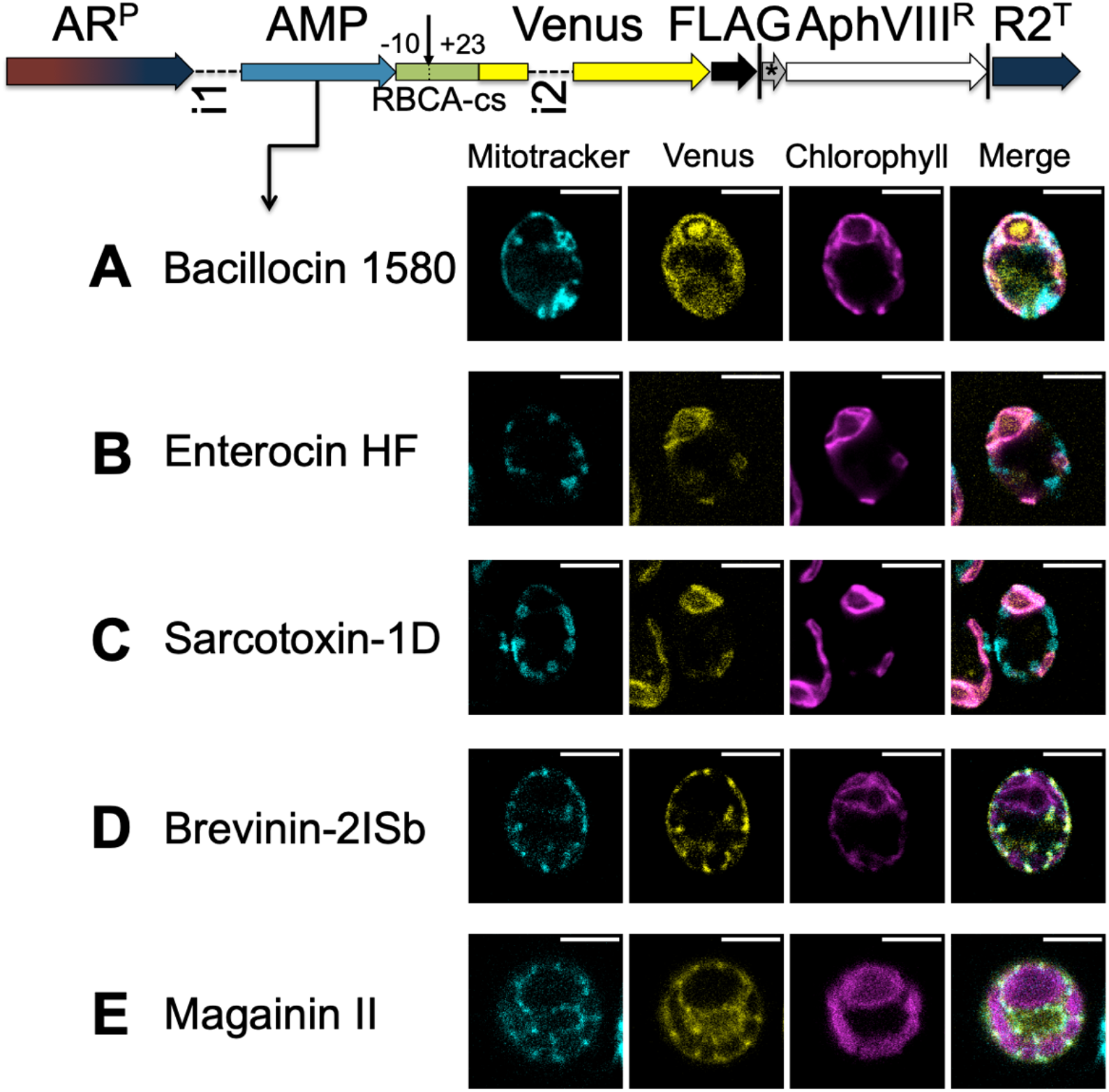
AMPs function as TPs. Constructs, schematically depicted at the top of the figure, assay the targeting ability of candidate peptides fused to the Venus-FLAG reporter, driven by the chimeric HSP70-RBCS promoter and the *RBCS2* 5’UTR (AR^P^) and RBCS2 terminator (R2^T^), and expressed bicistronically via the STOP-TAGCAT (*) sequence with the paromomycin resistance marker (*AphVIII^R^*). Vertical lines indicate stop codons. Expression levels in *C. reinhardtii* are increased by the use of introns: RBCS2 intron 1 (i1) in the 5’ UTR and RBCS2 intron 2 (i2) within the Venus coding sequence. Candidate HA-RAMPs, i.e. bacillocin 1580 (A), enterocin HF (B), sarcotoxin-1D (C), brevinin-2ISb (D) and magainin II (E) were fused to the RBCA cleavage site fragment encompassing residues −10 to +23 (RBCA-cs) and inserted upstream of Venus. The site of cleavage is indicated by a downward arrow. False-colour confocal images of representative cells show mitochondria as indicated by mitotracker fluorescence in cyan, the localisation of Venus in yellow and chlorophyll autofluorescence in magenta. Scale bars are 5μm. See Supplementary Figure 9 for a quantification of co-localisation, Supplementary Figure 10 for replicates, and Table S6 for a description of peptide sequences.

By contrast, when fused to two peptides with computationally generated random amino acid sequences followed by RBCA-cs, Venus fluorescence remained in the cytosol (Supplementary Figure 7B,C), showing that random peptides do not necessarily generate targeting in the presence of the RBCA-cs fragment. Furthermore, the class II HA-RAMP Brevinin 1E, fused to RBCA-cs, equally failed to deliver Venus into either organelle, appearing instead to accumulate in the vicinity of the chloroplast, particularly in one bright spot (Supplementary Figure 7D).

In order to demonstrate that the typical cells shown in Supplementary Figure 6 and Figure 5 are representative of the populations they were drawn from, we quantified co-localisation by calculating Pearson correlation coefficients (PPC) across fluorescence channels [59] for around 30 cells per strain (Supplementary Figure 9). Cells expressing the CAG2-mTP+, magainin II and brevinin-2ISb constructs had significantly higher PCCs between Venus and MitoTracker signals than cells expressing any other constructs, confirming mitochondrial localisation of Venus.

Similarly, the RBCA-cTP+, bacillocin 1580, sarcotoxin-1D and enterocin HF constructs gave rise to significantly higher PCCs between Venus and chlorophyll autofluorescence, indicating Venus does indeed reside in the chloroplast. Since nuclear transformation in *C. reinhardtii* results in random integration, we also checked that the genomic locus of integration did not influence targeting: import phenotypes were consistent across three independent insertion lines (Supplementary Figure 10).

For an independent assessment of Venus localisation, we isolated intact chloroplasts and mitochondria from whole cells of *C. reinhardtii* harbouring one chloroplast and one mitochondrial targeting construct (Figure 6). As previously described [60], chloroplast-enriched fractions still show some mitochondrial contamination due to the presence of a subpopulation of mitochondria firmly bound to chloroplasts in this microalga. In agreement with fluorescence imaging observations, bacillocin 1580-driven Venus-FLAG was absent from isolated mitochondria but present in whole-cell and chloroplast-fractions, just like the chloroplast markers OEE2 and RBCS (Figure 6A). Magainin II-driven Venus-FLAG behaved like mitochondrial markers COXIIb and F1β, being present in all three fractions (Figure 6B). Note that the mitochondrial fraction appears underloaded, likely due to an overestimation of protein concentration. Nonetheless, the strong FLAG signal in the whole cell fraction suggests that not all of the reporter protein is imported, in line with some Venus fluorescence originating from the cytosol in this strain (Figure 5E, Supplementary Figure 10K).

**Figure 6.**
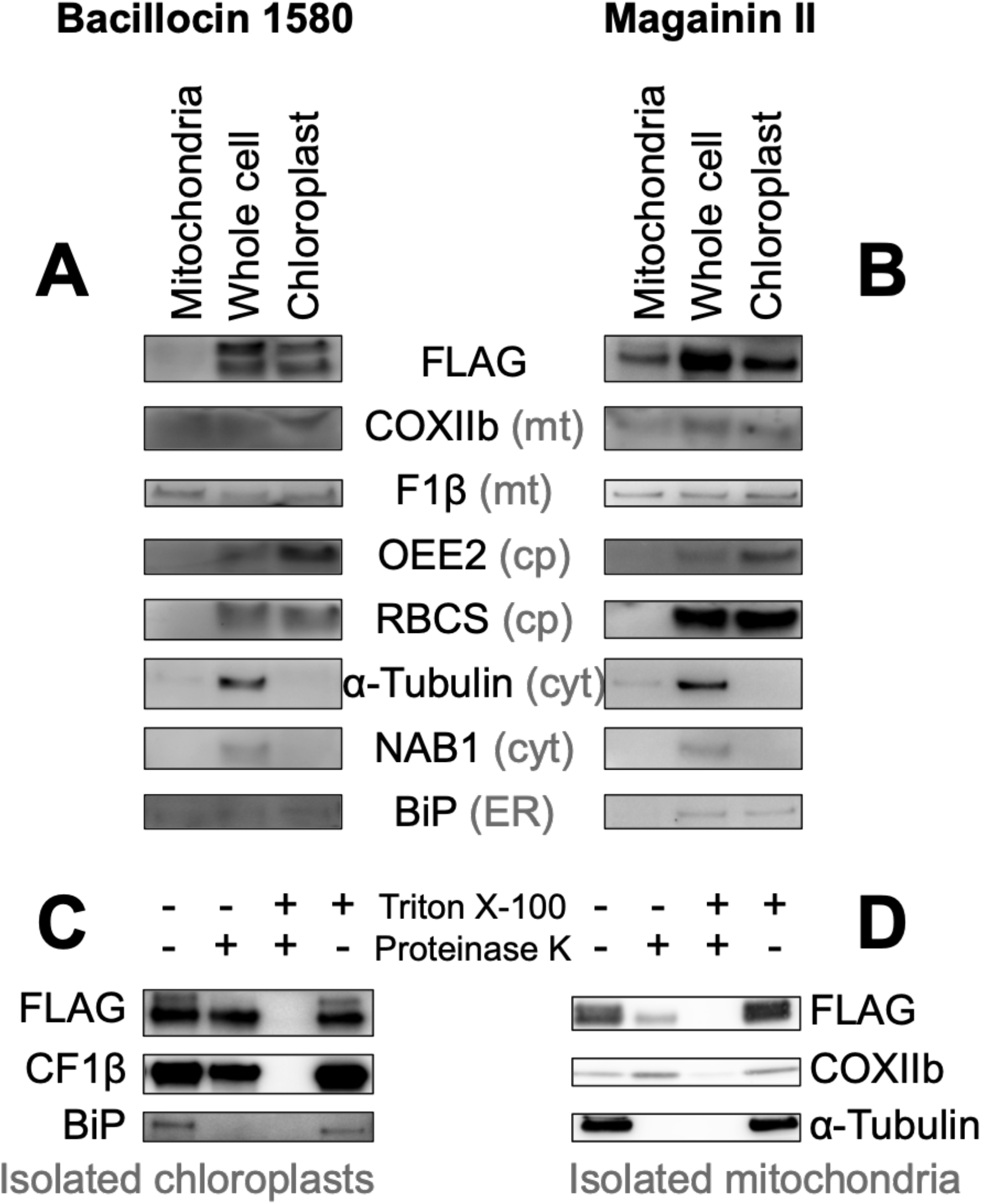
Biochemical confirmation of AMP targeting activity. Mitochondrial, whole cell and chloroplast fractions (1 *μ*g protein per well) isolated from Chlamydomonas strains in which Venus localisation is driven by bacillocin 1580 (A) or magainin II (B), each fused to the RBCA cleavage site, were immunolabelled with antibodies raised against against FLAG, an epitope tag carried C-terminally by the Venus reporter, and markers for different cellular compartments: Cytochrome Oxidase subunit IIb (COXIIb) and ATPsynthase subunit F1β for mitochondria (mt), Photosystem II Oxygen Evolving Enhancer 2 (OEE2) and Rubisco small subunit (RBCS) for chloroplasts (cp), α-Tubulin and nucleic-acid binding protein 1 (NAB1) for the cytosol (cyt) and lumenal binding protein (BiP) for the endoplasmic reticulum (ER). Isolated chloroplasts from the Bacillocin 1580 strain (C) and isolated mitochondria from the Magainin II strain (D) were subjected to a proteinase assay, where aliquots were treated with either 150 *μ*g ml^-1^ proteinase K and/or 1% Triton X-100, a membrane solubilising detergent. Aliquots were subsequently immuno-labelled with antibodies against FLAG, chloroplast ATP synthase subunit CF1 *β* and other organelle-markers described aside.

For both constructs, some of the Venus-FLAG reporter was protected from degradation by proteinase K in isolated organelles unless treated with detergents, again mirroring the behaviour of organelle-specific controls (Figure 6C, D). This confirms that bacillocin 1580 and magainin II act as *bona fide* TPs, with a significant fraction of the cargo protein localised inside the respective targeted organelle. A larger-sized fraction of Venus-FLAG did show sensitivity to proteinase K in the absence of detergents. This sensitivity mirrored that of tubulin and BIP, both minor contaminants in mitochondrial and chloroplast fractions respectively that are not protected within organelles and digested readily irrespective of the presence of a detergent. Thus, a subpopulation of AMP-reporter pre-proteins remains associated with the outer membrane of either organelle, likely as a result of incomplete or aborted translocation.

#### TPs show antimicrobial activity

To determine the AMP-activity of TPs, we performed the symmetrical experiment. Several chemically synthesised TPs, chosen for their variable proximity to class I HA-RAMPs (Supplementary Figure 8B), were used to challenge *Bacillus subtilis* in a standard assay [65] using magainin II as a positive control (Figure 7). *B. subtilis* was chosen as target organism rather than *E. coli,* the other standard laboratory bacterial species, because it proved more sensitive to HA-RAMP activity (Supplementary Figure 11). A high sensitivity was deemed a useful feature for an antimicrobial assay in this proof-of-principle experiment, since we expected TPs to have a lower activity than *bona fide* AMPs as they have been selected for targeting and not for impeding microbial growth over the last 1.5 By. All four tested *C. reinhardtii* TPs showed antimicrobial activity, as did F1β-mTP, the mTP of a mitochondrial ATP synthase subunit from *Neurospora crassa*, whose antimicrobial activity had previously been reported [76]. By contrast, neither the *A. thaliana* TL16-tSP, which targets proteins to the thylakoid lumen, nor the small hormone peptide cholecystokinin-22 (cck-22), here used as negative controls, inhibited growth demonstrating that antimicrobial activity is not simply an inherent feature shared by all peptides.

**Figure 7.**
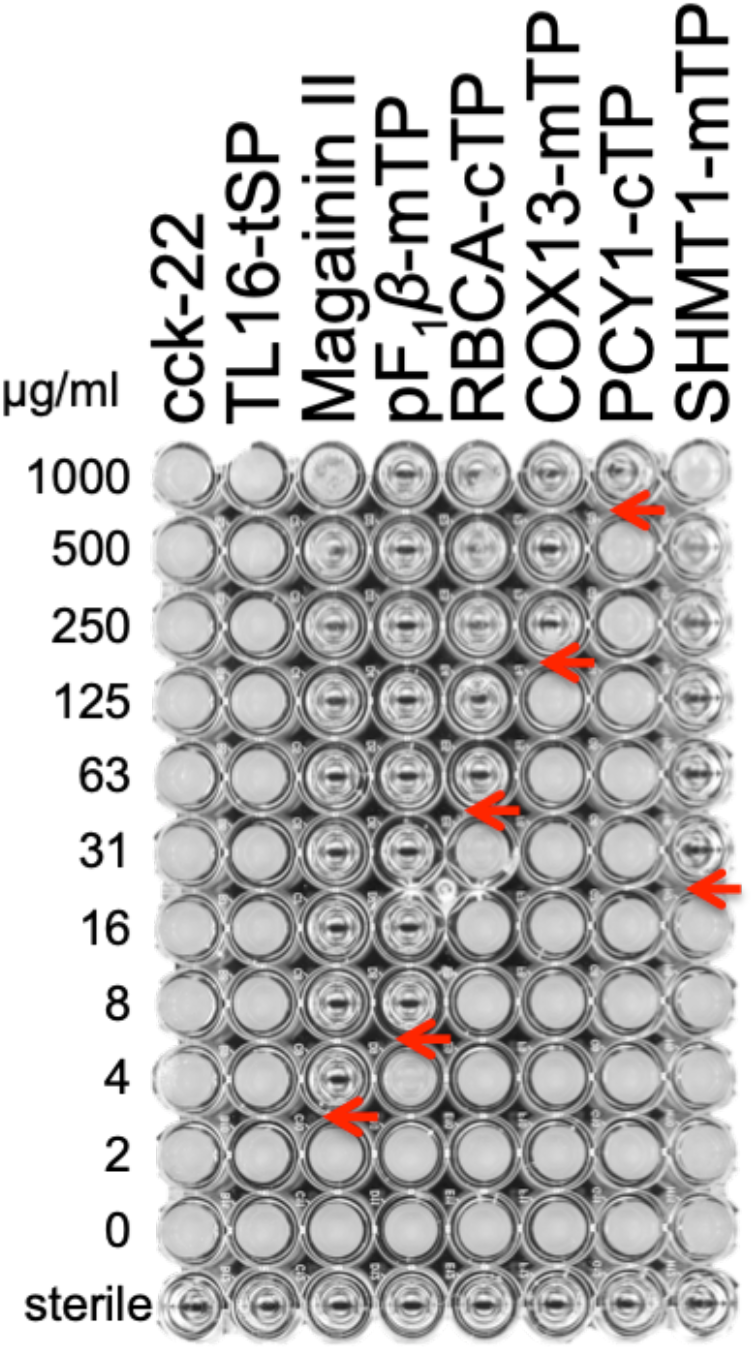
TPs exhibit antimicrobial activity. *B. subtilis* was challenged with serial dilutions of synthetic peptides (See Table S7 for sequences): the *Rattus norvegicus* peptide hormone cholecystokinin-22, the *C. reinhardtii* tSP of lumenal 16.5 kDa protein, the HA-RAMP magainin II, *Neurospora crassa* mTP of ATP synthase F1β subunit and *C. reinhardtii* cTPs from RBCA and plastocyanin (PCY1) or mTPs from cytochrome c oxidase 12 kDa subunit (COX13) and Serine HydroxylMethyl Transferase 1 (SHMT1). Transparent wells illustrate absence of growth. Red arrows point to the minimal peptide inhibiting concentration.

## Discussion

### Diversity of HA-RAMPs

In our *in silico* analysis, we first showed that HA-RAMPs can be grouped into distinct subtypes that do not always line up with the classification into antimicrobial families described in the literature (see Supplementary Text). Our analysis indicates that a systematic classification according to physico-chemical properties would be possible upon further investigation, which should prove of interest to the AMP community. However, it must be kept in mind that many bipartitions along the NJ clustering tree still have limited support.

### Evidence for a common origin of TPs with a class of HA-RAMPs

Our *in silico* and *in vivo* data support a common evolutionary origin of TPs and HA-RAMPs [11] as they have similar physico-chemical properties and show cross-functionalities.

Whether by k-means clustering (whatever the k-values) or by PCA analysis on the three first components, TPs consistently grouped together with class I HA-RAMPs and away from the three types of SPs targeting to bacterial periplasm, ER or thylakoid compartments, further emphasising their extensive similarities. These shared physico-chemical properties are consistent with an evolutionary link between a large subset of class I HA-RAMPs and TPs. On the other hand, the grouping of class II HA-RAMPs and SPs –-being both more hydrophobic than TPs and class I HA-RAMPS – is not robust in PCA and k-means analysis.

It had been argued that a fraction of random sequences (between 20 and 30% of those that were tested) could function as mitochondrial or secretory targeting peptides [77–79]. The random sequences that allowed functional targeting were strongly biased in sequence, with a requirement for a positively charged amphiphilic helix for proper interaction with the membrane surface [33]. This is in line with our own observation of an overlap between some random peptides, class I HA-RAMPs and TPs in our PCA analysis and with the shared properties of class I HA-RAMPs and TPs.

While the presence of amphiphilic helices in mTPs is well-established [31,32], the amphiphilic nature of cTPs had been questioned [37]. The present study shows that cTPs do display amphiphilic stretches capable of folding into amphiphilic helices, in line with NMR studies on selected cTPs in membrane-mimetic environments [38–40]. These helices are of similar length as those of mTPs, but make up a shorter proportion of the peptide in longer cTPs. Their amphiphilic character of cTPs may have been overlooked because the amphiphilic helix, which covers most of the shorter mTP sequences, is surrounded by additional elements, which do not fold into amphiphilic helices, such as an uncharged N-terminus [6,34–36] and a C-terminus with β-sheet characteristics [5,6].

Beyond similarities in physico-chemical properties, the proposed evolutionary relationship between TPs and HA-RAMPs was experimentally supported here by the antimicrobial activity observed for all tested TPs (Figure 7). It is very remarkable that TPs still display an antimicrobial activity, since they have not been selected for this function for the last 1.5 By [2]. It is not a surprise then, that higher concentrations of TPs, relative to the *bona fide* AMP Magainin II, are required to impede bacterial growth.

Further support for an evolutionary relationhip between these peptides stems from the organelle targeting abilities of the five HA-RAMPs we probed experimentally, such as bacillocin 1580, targeting the chloroplast, and magainin II, targeting the mitochondria (Figures 5 and 6). Our experiments using HA-RAMP-and TP-driven targeting to organelles, argue for a similar import process for both types of peptide through the canonical translocation pathways for mitochondria (TOM/TIM for Translocase of the Outer/Inner Membrane) and chloroplast (TOC/TIC for translocon on the outer/inner chloroplast membrane). Indeed, the Venus reporter is localised in the stroma or matrix with a post-import cleavage of the N-terminal pre-sequences. By contrast, non-canonical targeting, which has been identified in a very limited number of cases [80,81], including a handful of glycoproteins [82], involves proteins that lack a cleavable pre-sequence and are delivered to envelope compartments -outer or inner membrane, or inter-membrane space – but not to the organelle interior [5,7,83].

One could argue that, rather than the present evolutionary scenario, a convergent evolution of class I HA-RAMPs and TPs driven by strong selective constraints could have led these peptides to adopt the same optimum in their physico-chemical properties. However, the respective antimicrobial and intracellular targeting functions of class I HA-RAMPs and TPs do not *per se* constitute a selective pressure for convergent evolution: as documented in the present study, cyclotides globular AMPs, as well as class II HA-RAMPs are clearly distinct from class I HA-RAMPs despite a shared antimicrobial function. Similarly, SPs function as cleavable N-terminal targeting sequences like TPs but do not group with class I HA-RAMPs nor do they show antimicrobial activity.

### Efficient targeting to extant organelles requires specific sequences besides the amphiphilic helix of TPs

Previous studies showed that short cTPs rely on the N-terminal region of the mature protein to allow chloroplast targeting [73]. Accordingly, cTPs being shorter in Chlamydomonas than in plants [this study and [71]], post-cleavage site residues are critical for proper chloroplast-targeting in this alga, as we demonstrated here for RBCA. This prompted us to include RBCA post-cleavage site residues in our HA-RAMP constructs. The mechanistic contribution of these mature N-termini is still unclear, but they could provide an unfolded stretch long enough to elicit import [73].

A major issue in targeting to intracellular organelles in plants and algae is the ability of a given presequence to avoid dual targeting. Several in vitro studies suggest that specific targeting is achieved, at least in part, through competition between the two organelle import systems. For instance, isolated mitochondria import cTPs [84,85] whereas, non-plants mTPs can drive import into isolated chloroplasts [86,87]. To avoid miss-targeting, plant and algal TPs have probably further evolved some specific traits of the N-terminal peptide region for targeting to chloroplasts [34,88,89]. Targeting specificity has been improved further with the acquisition by mTPs of a chloroplast avoidance signal consisting in multiple Arg at their N-terminus [36]. In agreement with this proposal, we note that, among the chloroplast-targeting HA-RAMPs, Bacillocin 1580 carries no charge within the first ten residues, and Enterocin HF carries a single Lys only. However, Sarcotoxin 1D has four charged residues within this N-terminal window, including two Arginines which might have been expected to exclude the construct from the chloroplast [36].

Other cTP motifs have been suggested to play some role in chloroplast protein import, such as Hsp70 binding sites within the first 10 residues of the peptide [35,55], or FGLK motifs, grouping aromatic (F), helix-breaking (G), small hydrophobic (L) and basic (K) residues for interaction with TOC receptors [35,56]. Bacillocin 1580 indeed contains one FGLK-site (sensu [56]), but neither enterocin HF, sarcotoxin-1D nor the RBCA presequence do, while the mitochondrial-targeting magainin II contains two. Clearly, further work is needed to understand mitochondrial versus chloroplast targeting for the HA-RAMPs under study.

### Targeting peptides and the translocation machinery

Some bacterial Omp85 outer membrane assembly factors, which target proteins by a C-terminal phenylalanine [90] and are thought to have given rise to TOC75, a core component of TOC [91]. Since Rhodophyte and Glaucophyte cTPs start with a conserved phenylalanine [92], chloroplast protein import could have benefited from a functional inversion of this cyanobacterial protein in the evolution of the TOC complex. Although this observation was taken as an argument against the emergence of cTPs from HA-RAMPs [93], we argue that HA-RAMPs would have originally interacted with the outer membrane lipid headgroups [5], then crossed the outer membrane spontaneously [94,95], with some HA-RAMPs also interacting with Omp85 proteins [96]. In this view, the most likely evolutionary scenario for chloroplast import has involved recruitment of Omp85 to improve delivery of HA-RAMP-tagged proteins to the import-and-destroy receptor at the inner membrane surface.

The bacterial resistance apparatus at the origin of the chloroplast protein translocon, aimed to prevent the lethal disruption of plasma membrane integrity by AMPs [22,23] is most likely to be found in the TIC rather than the TOC part of the translocon. It should be emphasized that cTPs have evolved in a context widely different from that prevailing for the emergence of mTPs. The latter indeed appeared in absence of any pre-existing import system in the archeal ancestor of eukaryotic cells. In contrast, the eukaryotic ancestor of Archeplastidia was in some way “preadapted” for the recruitment of Class I HA-RAMP for import functions. This does not mean that cTPs merely have recruited the TOM/TIM that form the protein channel as well as Tic21, Tic22, Tic23, Tic32, Tic55 and Tic62 are of cyanobacterial origin [97–99]. However, we anticipate a common origin of some TIC and TIM subunits which will require an extensive phylogenetic analysis of the two sets of translocon components.

### Conclusion

Although evolutionary scenarios necessarily give rise to conflicting views, it should be kept in mind that neither the scenario of convergent evolution nor that of a spontaneous generation of TPs account for the emergence of the ancestral mitochondrial and chloroplast translocation systems. In contrast, an antimicrobial origin of TPs is a more parsimonious scenario, in which the import-and-destroy ancestral mechanism allowing the endosymbiont to resist the attacks of AMPs [20–26] is at the root of the translocation systems [11]. Further support for this view came from recent studies of the amoeba *Paulinella chromatophora,* which acquired a novel primary endosymbiotic organelle called “chromatophore” approximately 100 million years ago [100]. Proteomic analysis of these chromatophores identified a large set of imported AMP-like peptides as well as chromatophore-imported proteins harbouring common N-terminal sequences containing AMP-like motifs [101]. These findings thus provide an independent example of a third primary endosymbiosis that is accompanied by the evolution of an AMP-derived protein import process. The detailed evolutionary histories of extant organelle translocons and bacterial transmembrane channels involved in AMP-resistance mechanisms should provide a means to further assess the antimicrobial origin of organelle-targeting peptides.

## Author Contributions

Conceptualisation: FAW, IL; Data Curation: CG, ODC, IL; Funding Acquisition: YC, FAW, IL; Investigation and Validation: CG and IL (computational), ODC (experimental); Software: CG, ODC; Supervision: YC, FAW, IL; All authors contributed to writing and editing the manuscript.

## Acknowledgments

We thank Richard Kuras, Ulrike Endesfelder, Michael Schroda and Moritz Meyer for fruitful discussions and advice, Masayuki Onishi and the Pringle lab, for sharing their bi-cistronic expression plasmids, Pierre Crozet for advice on plasmid design, Zhou Xu and Simon Desjardin for help with microscopy, and Barry Bruce and Prakitchai Chotewutmontri for their help in the correct implementation of Hsp70 binding site and FGLK motif search scripts. This work was supported by the annual funding from the Centre National de la Recherche Scientifique and Sorbonne University to UMR 7141, by the ChloroMitoRAMP ANR grant (ANR-19-CE13-0009) and by LabEx Dynamo (ANR-LABX-011). ODC was supported by The Rothschild Foundation, the Labex dynamo and the ChloroMitoRAMP grant. CG was supported by the MATHTEST ANR grant (ANR-18-CE13-0027).

