## Supplementary Material for "Evidence supporting an antimicrobial origin of targeting peptides to endosymbiotic organelles"

### Supplementary Text

*Diversity of HA-RAMPs*

Rather than reflecting structural relationships between peptides, AMP families have often been named after the source of novel peptides or a defining characteristic. For example, gaegurins, named after the Korean word "Gaegury" for frog, regroup AMPs isolated from the Korean frog *Glandirana emeljanovi* (formerly *Rana rugosa*) even though they are closer in sequence to other HA-RAMP families than to each other [102]. Unsurprisingly, gaegurins are spread across both HA-RAMP classes (Figure 1C). However, not all idiosyncratically named families lack structural systematics. As more and more AMPs were being characterised, investigators started to class novel peptides that were deemed related in primary sequence in families named after the first described peptide of that type. Cecropins, named after the giant silk moth *Hyalopphora cecropia* from which the first members were extracted, are now a well-established family including sarcotoxin or bactericidin peptides. They form a relatively uniform group to the right of the NJ clustering tree (Supplementary Figure 2). Conversely, brevinin-1 and brevinin-2 carry the same name because they were originally isolated together from the Japanese frog *Pelohylax porosus* (formerly *Rana brevidopda porsa*), but are now two separate families, each expanded by addition of homologous peptides [103]. The brevinin-1 family forms a fairly homogeneous branch towards the bottom of the clustering tree (lower left side on Supplementary Figure 2), interspersed only by some gaegurin and ranatuerin peptides that had already been recognised as brevinin-1 homologues [102,103]. Brevinin-2, by contrast, is split into three relatively distinct subgroups in our analysis. It is interesting to note that brevinin, esculentin and ranatuerin families all containing the "Rana box" span both class I and class II HA-RAMPs (Figure 2C). The "Rana box" is a C-terminal disulphide bridge motif shared among ranid AMPs [104] that suggests a common origin and shows that related HA-RAMPs diversify rapidly in terms of physico-chemical properties.

### Supplementary Figures

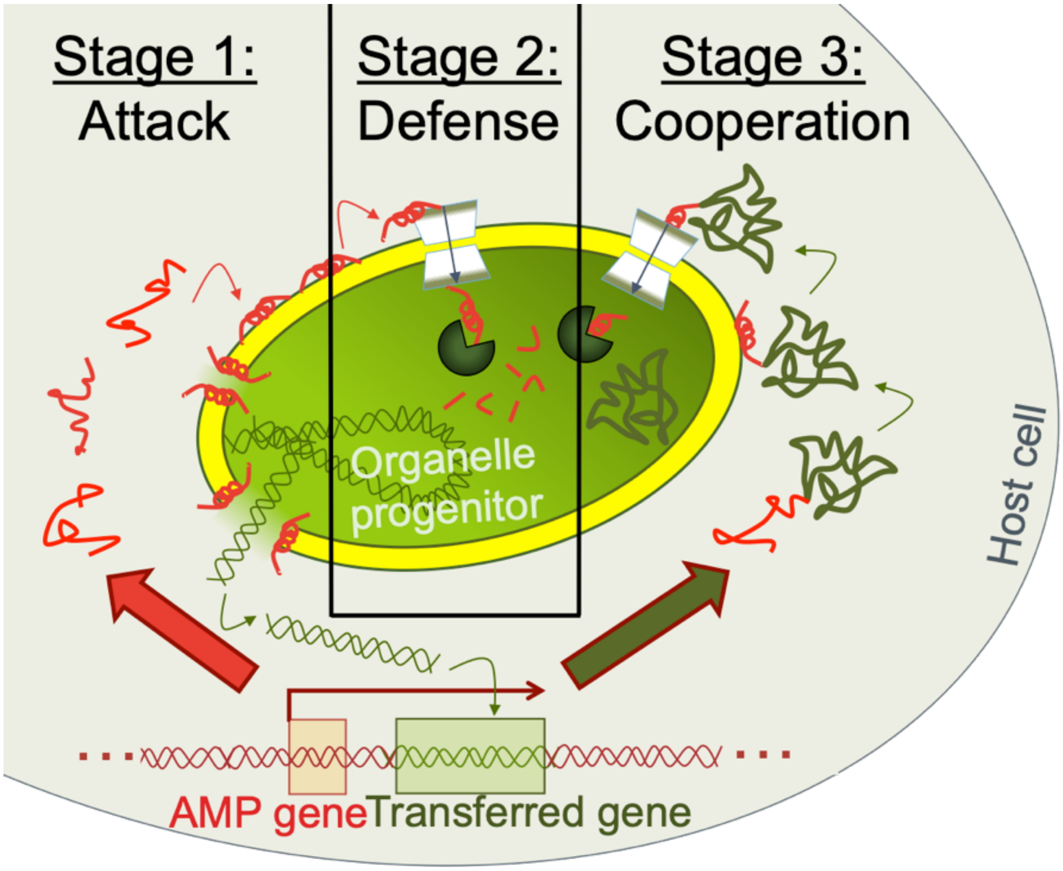


**Supplementary Figure 1. Emergence of TPs from AMPs implies a three-stage scenario for the evolution of endosymbiotic protein targeting systems.** In stage 1, the host attacks the proto-endosymbiont using ribosomally synthesised AMPs. Cell lysis releases genetic material, which may occasionally be integrated into the host genome. In stage 2, the proto-endosymbiont acquires an import-and-destroy mechanism to resist host attacks, comprising a dedicated AMP transporter and a cytosolic peptidase. In stage 3, this detoxification mechanism is co-opted to import any protein that results from serendipitous fusion of genes downstream of AMP coding sequences.

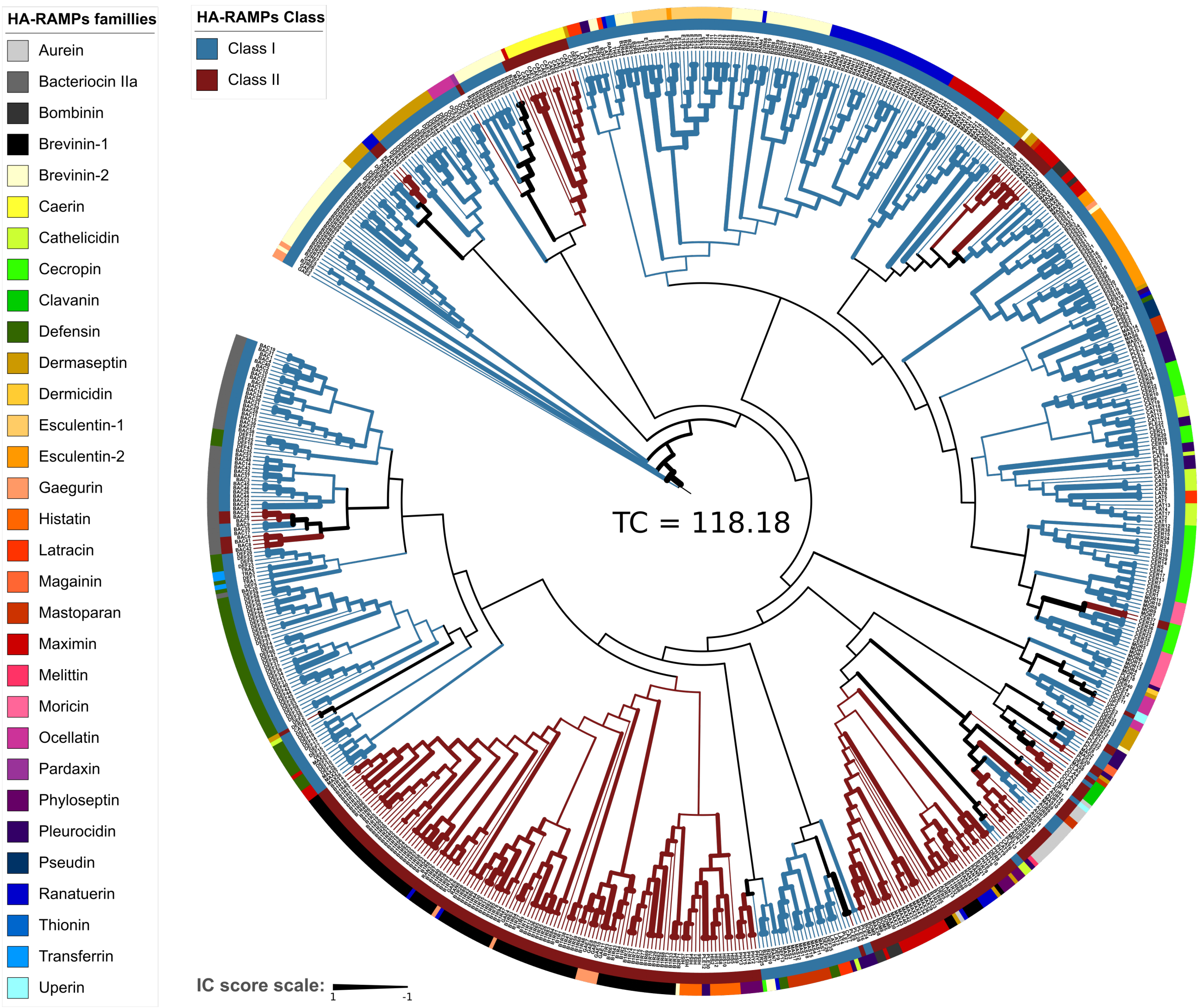


**Supplementary Figure 2**. **HA-RAMPs families and classes are spread all over the tree.** Neighbour-joining tree based on Euclidean distances of HA-RAMPs described by 36 ACC terms. The branches are coloured according to HA-RAMP classes I (blue) and II (red), as defined by the k-means clustering (Figure 1). Branch widths are proportional to the Internode Certainty (IC) value, indicating the robustness of the tree (see Methods). IC values at or close to −1 (thin branches) indicate an almost complete absence of support for the bipartition defined by the branch among bootstrap trees and IC values close to 1 (thick branches) indicate the absence of conflict among the bootstrap trees [52]. The sum of IC values for all branches (Tree certainty, TC) is given at the centre of the tree. When all children of a node have the same colour, the colour propagates inwards towards the root. The inner circle around the NJ tree indicates the HA-RAMP class of the peptide. The outer circle indicates the family of the peptide, as described in the literature. See figure box for colour code. Class I HA-RAMPs are found in different parts of the NJ clustering tree, sometimes together with some class II HA-RAMPs (in red), but the two classes are not intermingled and tend to form robust homogeneous sub-trees.

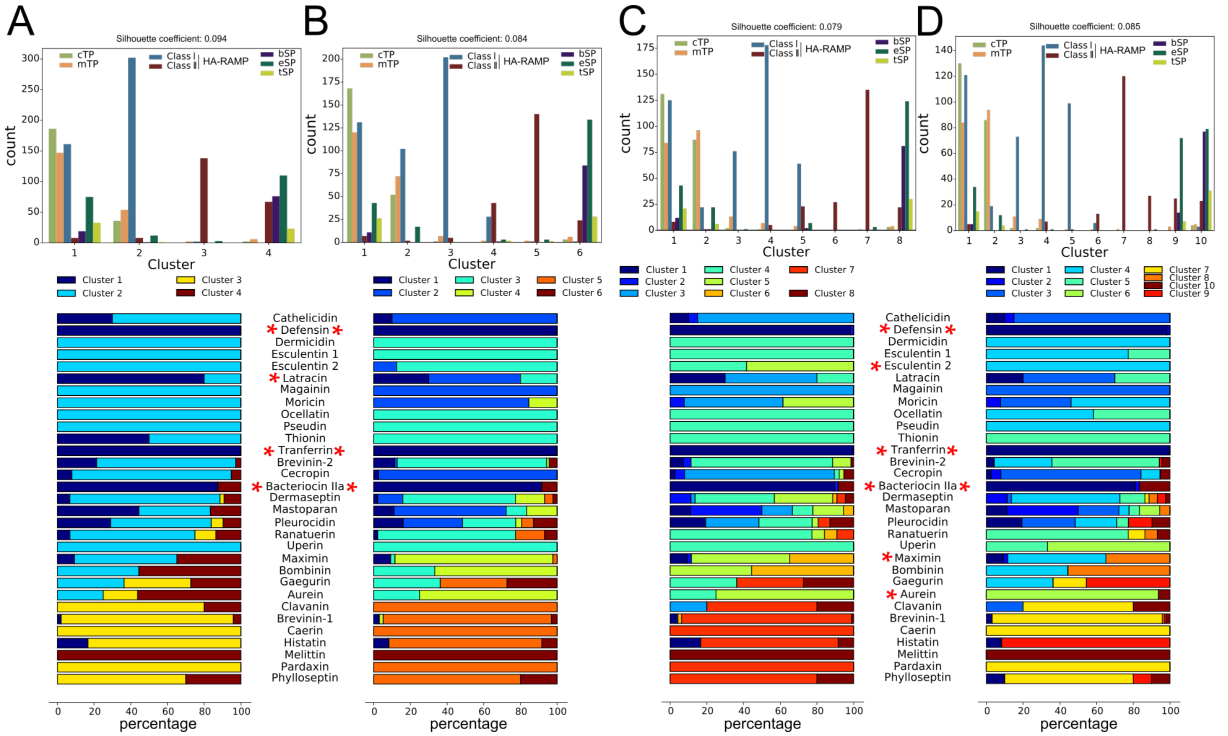


**Supplementary Figure 3**. **TPs cluster with class I HA-RAMPs.** K-means clustering of peptides, as described by their 36 ACC terms, with cTP (green), mTP (orange), class I HA-RAMP (blue), classII HA-RAMP (dark red), bSP (dark green) and tSP (light green) ($Chi^2$ Pearson test, $p < 4.94 \cdot 10^{-324}$). From (A) to (D), k-means clustering with k= 4, 6, 8 and 10. Top: The average silhouette coefficient is indicated on the top of the cluster distribution. The cluster at the most left side contains a majority of TP and class I HA-RAMPs. This cluster is robust and conserves about 60% of its peptides from k=3 to k=10. Bottom: percentage of peptides of HA-RAMP families described in the literature among the k-means clusters. Families that are mainly found in the robust cluster (at the most left side in the first column) are indicated by a red star.

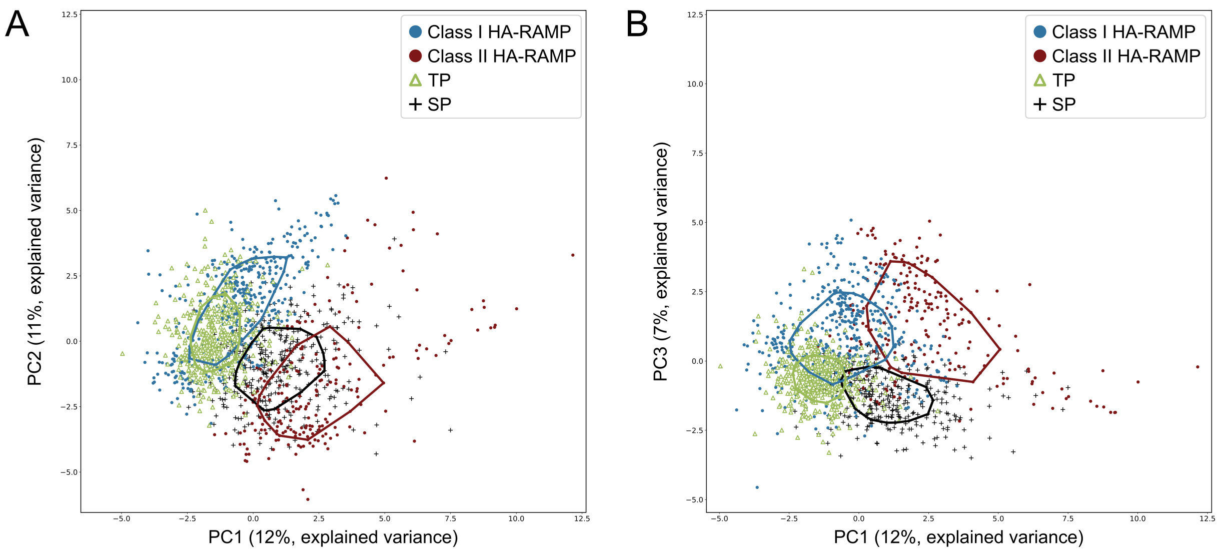


**Supplementary Figure 4. The third principal component of the PCA analysis confirms proximity between class I HA-RAMPs and TPs, while improving the separation between class II HA-RAMPs and SPs**. (A) PCA on normalised ACC terms for class I HA-RAMP (blue circle), class II HA-RAMP (dark red circle), targeting peptides (green triangle) and signal peptides (black cross). Peptide positions are plotted along the first (X) and second (Y) principal components. Note that the z1.lag1 ACC term reflecting the hydrophobicity has the largest contribution in PC2 (Supplementary Figure 5C). (B) Same PCA as in (A) with peptide positions plotted along the first (X) and third (Y) principal components. Note that z1.lag4 reflecting the amphiphilic character of the helix has the largest contribution in PC3 (Supplementary Figure 5C). Explained variance is indicated in parenthesis for each axis. Solid lines represent the convex areas of the 50% most central peptides in each group. ACC term contribution in the principal components are given in Supplementary Figure 5 C,D.

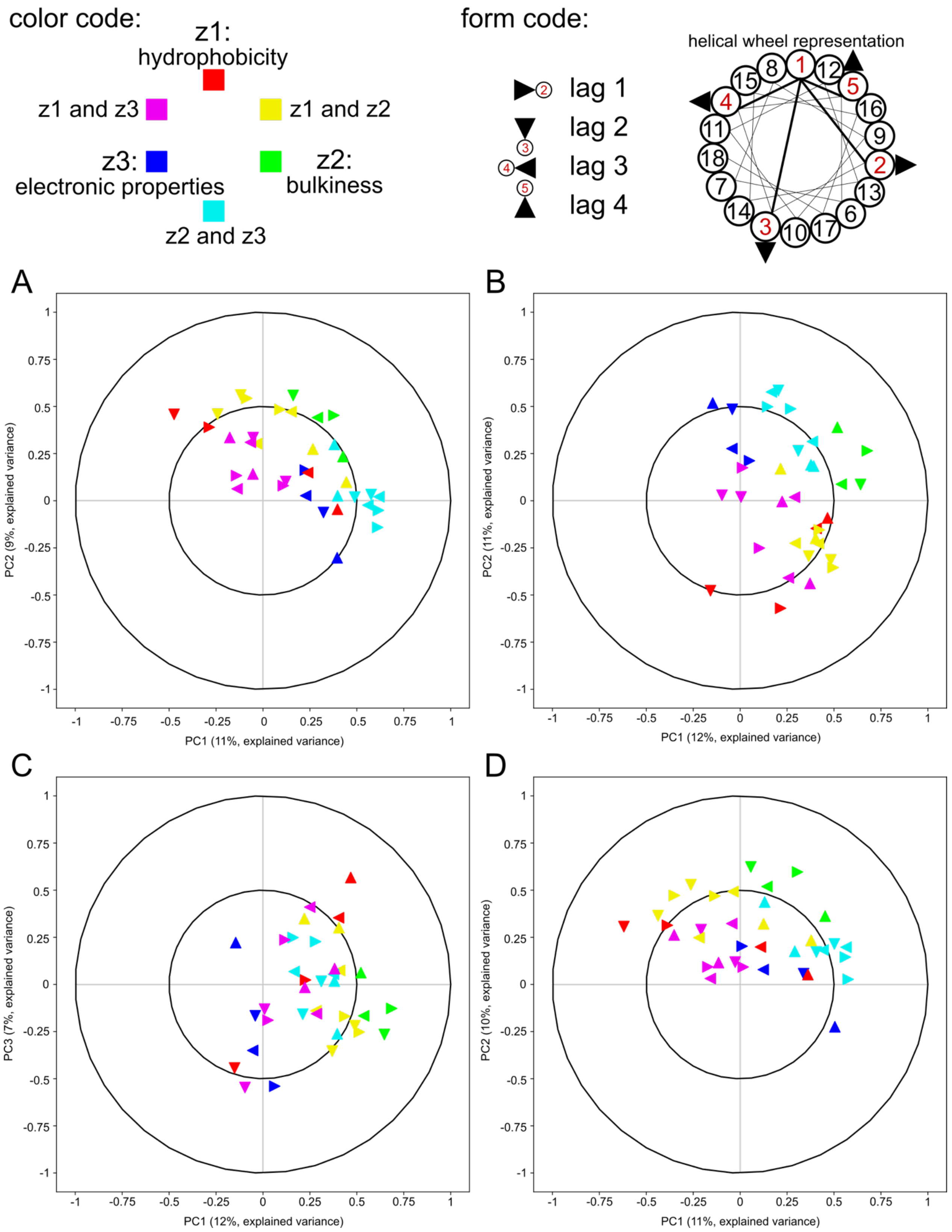


**Supplementary Figure 5. Separation of peptides in PCA analyses reflects hydrophobic and**

**amphiphilic features**. (Top scheme) Each triangle represents an ACC term, where the colour corresponds to the z-scales involved as described in the legend, and the orientation corresponds to the lag as indicated on the helical wheel representation: a lag of 1 thus describes the relation between residue 1 and residue 2; a lag of 2 between residue 1 and residue 3 etc. The maximum considered is lag 4, thus only residues 1-5 are concerned. The red numbers in the circles next to the triangles indicate the residues considered relative to the first one, highlighted in red on the helical wheel representation. For a qualitative analysis of the physico-chemical significance of these two components, one has to keep in mind that an α-helix being 3.6 amino acids per turn, the n+3 and n+4 residues (reflected by lag 3 and lag 4 terms) can be considered as lying on the same face of the α-helix as residue n, whereas the n+1 and n+2 residues (reflected by a lag 1 and lag 2 terms) rather lie on the opposite face.

(A) Correlation circles for the PCA presented in Figure 4. The coupling between electronic and steric properties  (z2.3 and z3.2) of the residues from the two faces of the amphiphilic helix (lag1 and lag3) are the main contributors to PC1 (largest distance from the origin along the x-axis) whereas the hydrophobic and steric properties- and their coupling- (z1, z2, z1.2 and z2.1) of the residues along the same face of the helix (lag1 and lag2) most contribute to PC2 (largest distance from the origin along the y-axis). Note that z1.lag2 is the term with the highest distance from the radius of the circle and define a diameter line with z3.lag4, meaning that z3.lag4 and z1.lag2 are the most anti-correlated terms. (B) Correlation circles for the PCA presented in Supplementary Figure 4A. Note that the z1.lag1 term, reflecting the hydrophobicity on tbe opposite side of the helix has the largest contribution in PC2. (C) Correlation circles for the PCA presented in Supplementary Figure 4B. Note that z1.lag4 reflecting the hydrophobicity on the same side of the helix has the largest contribution in PC3. (D) Correlation circles for the PCA presented in Supplementary Figure 8. X-axis and y-axis are principal components used for representation. Unit circle and half unit circles are represented.

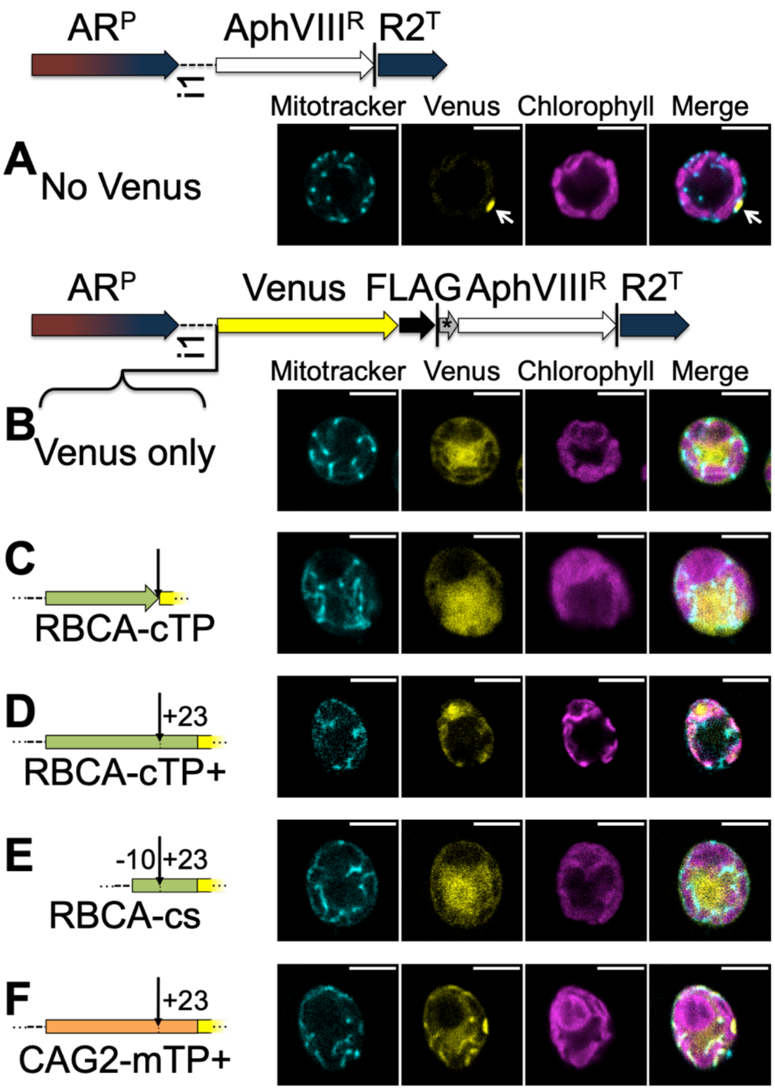


**Supplementary Figure 6. A RBCA cleavage site including downstream residues is necessary but not sufficient for targeting.** False-colour confocal images of representative C. reinhardtii cells show mitochondria as indicated by mitotracker fluorescence in cyan, the localisation of Venus in yellow and chlorophyll autofluorescence in magenta. Scale bars are 5$\mu$m. Expression constructs use the chimeric HSP70-RBCS2 promotor ($AR^P$), RBCS2 intron 1 (i1) in the RBCS2 5’ UTR, the paromomycin resistance gene ($AphVIII^R$) as selectable marker, and the RBCS2 terminator ($R2^T$). Vertical lines represent stop codons. A Venus fluorescent reporter carrying a FLAG-tag at the C-terminus was introduced upstream of $AphVIII^R$, and bicistronic expression ensured by connecting the two genes with a STOP-TAGCAT sequence (*). Candidate peptides to be assayed for targeting were then introduced upstream of Venus. (A) shows cells expressing the selectable marker only (the white arrow marks the eyespot). (B) Venus fluorescence is shown in the absence of a presequence, (C) when fused to Rubisco activase (RBCA) cTP up to the cleavage site, (D) including 23 residues downstream of the cleavage site, (E) the RBCA cleavage site (RBCA-cs) fragment alone encompassing residues -10 to +23, (F) or fused to $\gamma$-carbonic anhydrase 2 (CAG2) mTP. In each case, the site of cleavage is indicated by a downward arrow. See Supplementary Figure 9 for a quantification of co-localisation, Supplementary Figure 10 for replicates, and Table S6 for a description of peptide sequences.

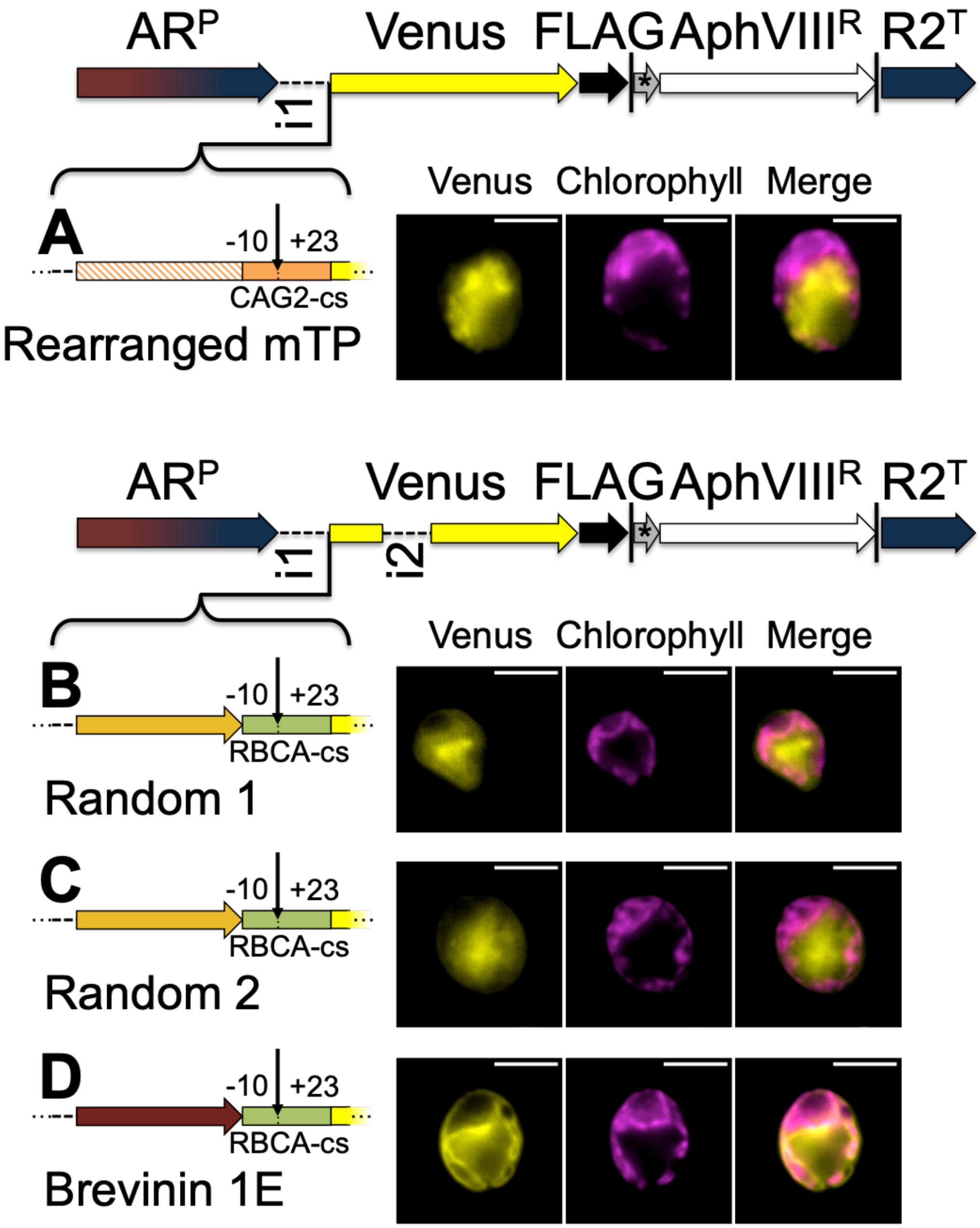


**Supplementary Figure 7. The presence of an amphiphilic helix is necessary but not sufficient for targeting.** False-colour epifluorescence images of representative C. reinhardtii cells show the localisation of Venus in yellow and chlorophyll autofluorescence in magenta. Scale bars are 5$\mu$m. See Supplementary Figure 8 for a description of construct elements; i2 is RBCS2 intron 2. (A) shows Venus fused to the CAG2-mTP, rearranged to be unable to form an amphipathic helix, and with residues (-10 to +23) encompassing the cleavage site (CAG2-cs). Lower panels show Venus localisation driven by computationally generated random peptides (B, C) or the class II HA-RAMP

Brevinin 1E, each fused to RBCA-cs. See Supplementary Figure 10 for replicates, and Table S6 for peptide sequences.

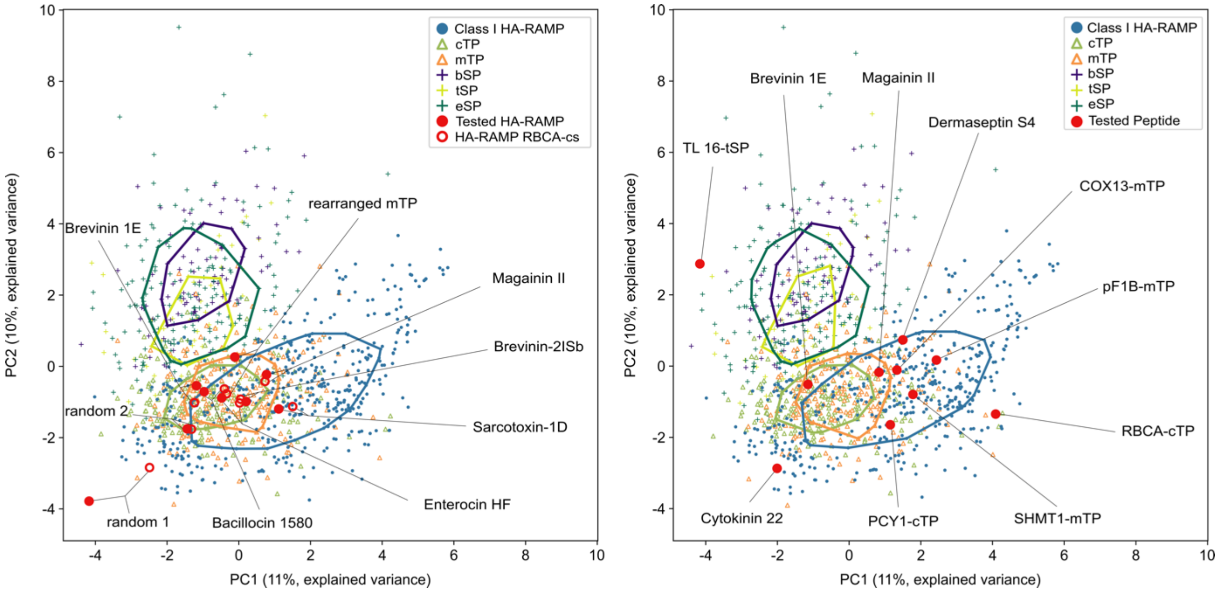


**Supplementary Figure 8. Position in PCA of class I HA-RAMPs and TPs selected for experimental analysis.** Same principle as in Figure 4, considering TP, class I HA-RAMPs and SP. (A) HA-RAMPs used for targeting assays are indicated (red dot: original, red circle: construct including the Rubisco activase cTP cleavage site, RBCA-cs). (B) Peptides used for antimicrobial assays are indicated in red. See Supplementary Figure 5 for the contribution of ‚ACC terms to the principal components PC1, PC2 and PC3 and Table S6 and S7 for peptide sequences.

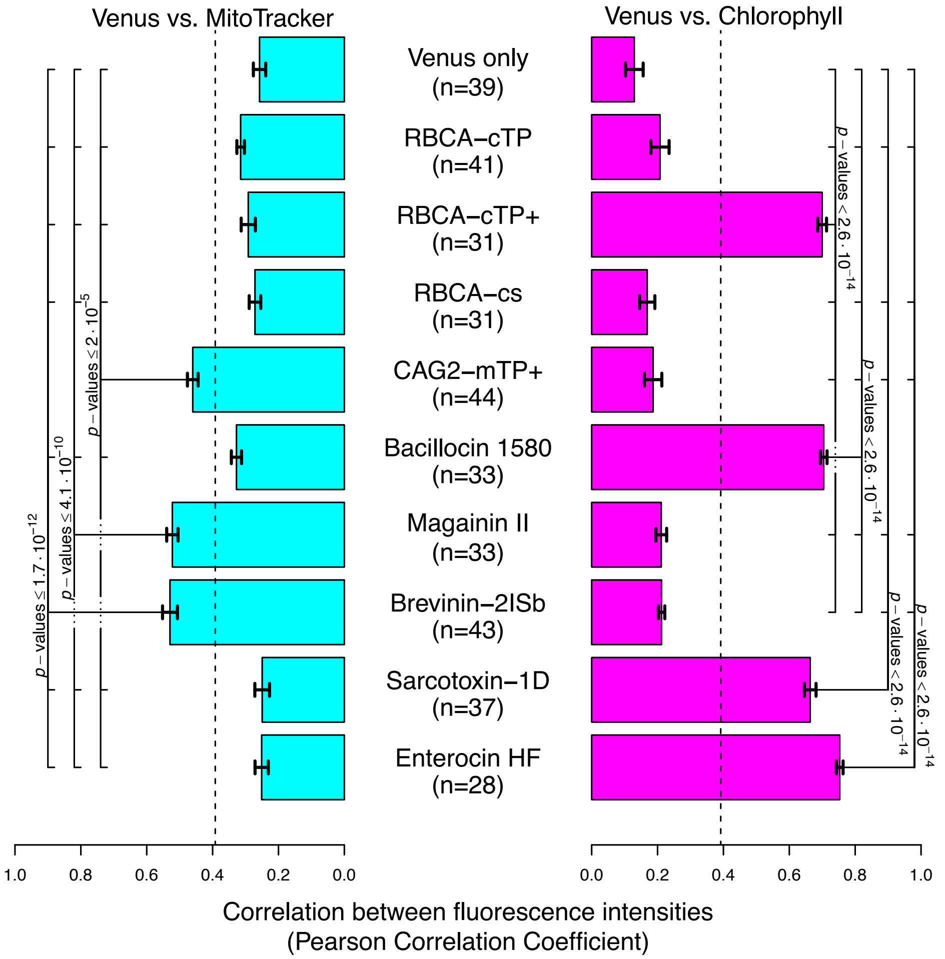


**Supplementary Figure 9. A quantitative assessment of co-localisation backs up targeting interpretations**. Pearson correlation coefficients were calculated for each strain between the Venus channel and either the MitoTracker or the chlorophyll channel of n cells were plotted as mean ± standard error. Perfect co-localisation would give a value of 1, whereas 0 would indicate completely uncorrelated localisation. The dashed line indicates the upper limit of a 95% confidence interval for PCCs between MitoTracker and chlorophyll channels across all strains, which serves as a control for random co-localisation that arises simply due to proximity of compartments within a cell. See Supplementary Figure 6 and Figure 5 for description of constructs and exemplary images. The “no Venus” strain was excluded from this analysis, as in the absence of Venus, Venus channel intensities within the cell are on the order of background values substracted from other strains (except for the eyespot). Significance was tested via a one-way ANOVA followed by a Tukey post-hoc analysis. All p-values <0.05 from the Tukey analysis are indicated.

171
172

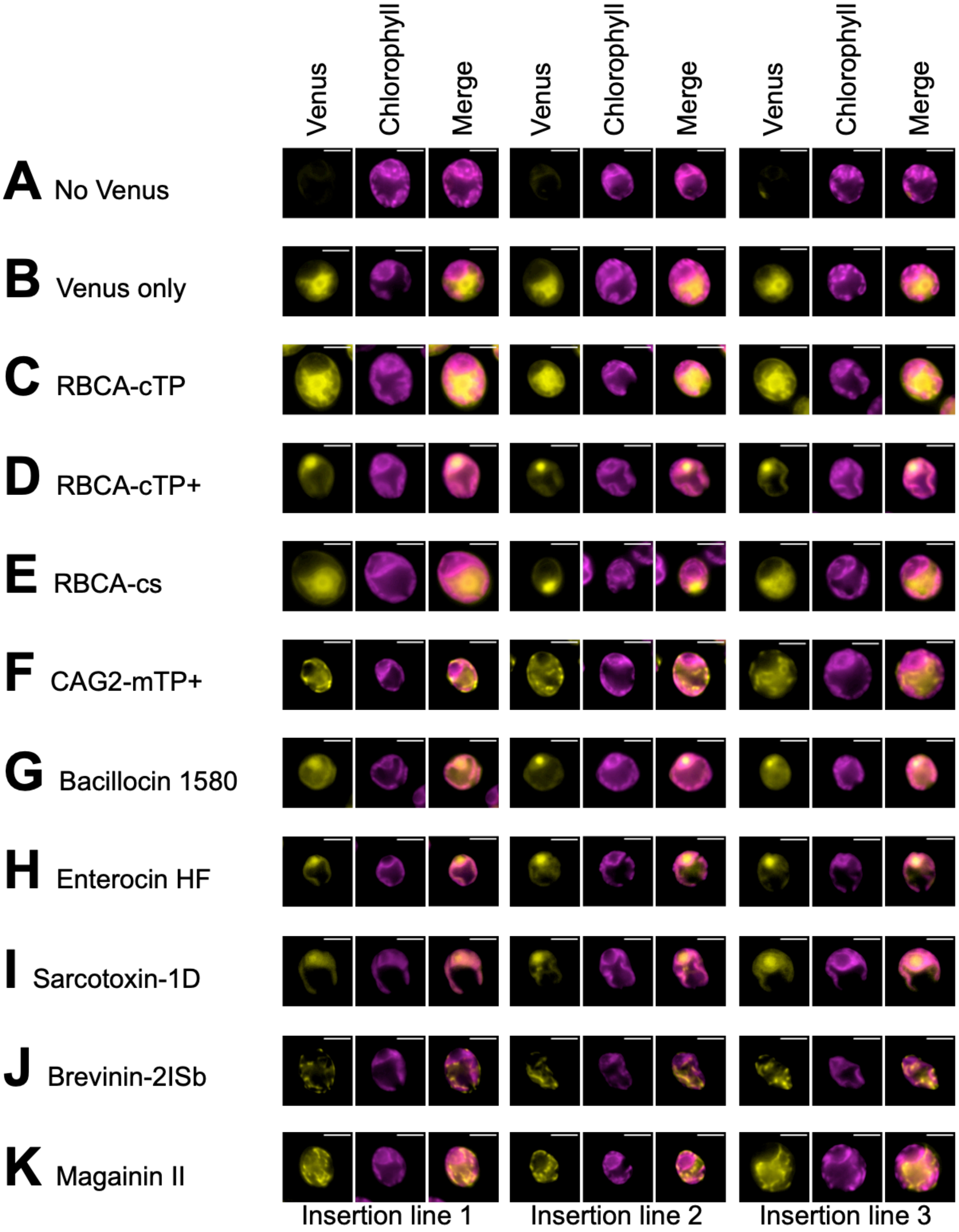


173

174 **Supplementary Figure 10. Three biological replicates all show the same phenotype for each**
175 **construct**. Epifluorescence false-colour images show representative cells from three independent

insertion lines for each construct. See Supplementary Figure 6 for a description of constructs (A) to (F), and Figure 5 for a description of constructs (G) to (K) and Supplementary Figure 7 for a description of constructs (L) to (O).

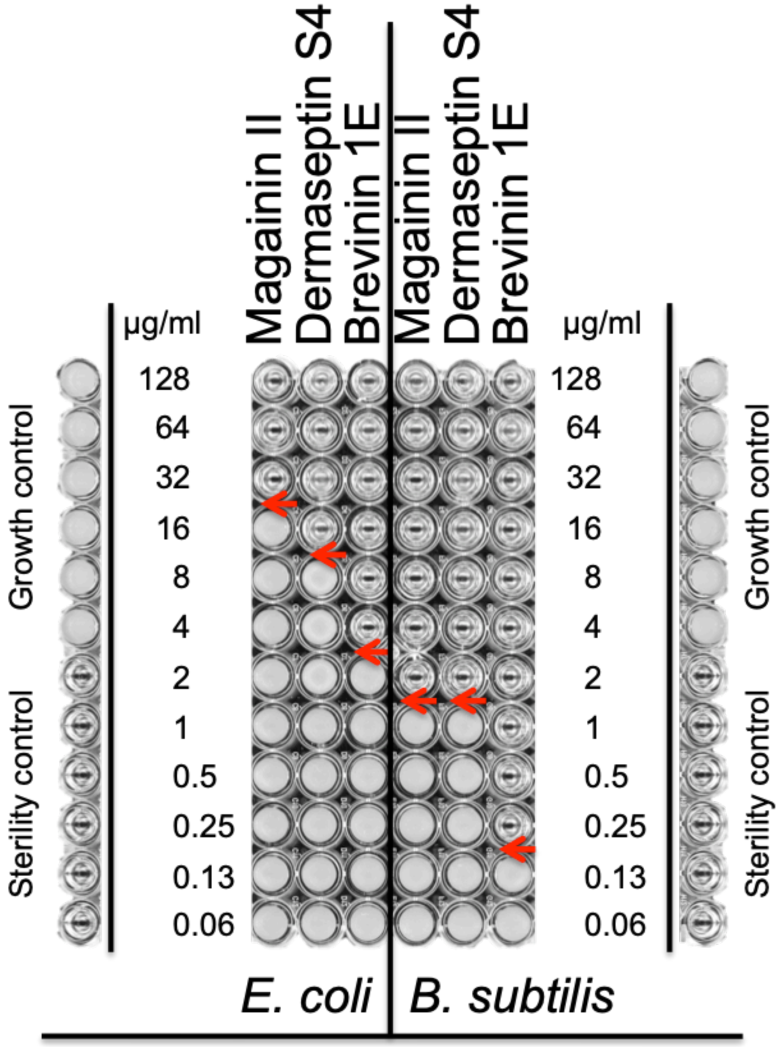


**Supplementary Figure 11. *B. subtilis* is a more sensitive probe for antimicrobial activity of peptides than *E. coli.*** Gram negative *E. coli* and gram positive *B. subtilis* were challenged with serial dilutions of the ynthetic HA-RAMPs Magainin II, Dermaseptin S4 and Brevinin 1E (See Table S7 for sequences). Transparent wells illustrate absence of growth. Red arrows point to the minimal peptide inhibiting concentration. All wells shown here are part of the same 96-well plate; control rows are shown separately on either side for improved clarity.
